# Impaired Calcium Signaling in Precapillary Sphincters and Pericytes Perturbs Neurovascular Regulation after an Ischemic Stroke

**DOI:** 10.64898/2026.01.11.698861

**Authors:** Lechan Tao, Chen He, Teddy Groves, Kayeon Kim, Krzysztof Kucharz, Aleksandra Petrovskaia, Dmitry Postnov, Xiao Zhang, Christina Latt Fjorbak, Henry G. Sansom, Hao Hu, Peter Andersen, Gavrielle R. Untracht, Inge A. Mulder, Ed Van Bavel, Martin Lauritzen, Anpan Han, Changsi Cai

## Abstract

Ischemic stroke disrupts neurovascular uncoupling, in which neuronal activity fails to evoke appropriate microvascular blood flow responses, despite successful recanalization of upstream arteries. The cellular mechanisms underlying this dysfunction remain poorly defined. We examined Ca²⁺ signaling and contractile dynamics of vascular smooth muscle cells, precapillary sphincters and contractile pericytes, using two-photon microscopy and laser speckle imaging in awake mice subjected to transient middle cerebral artery occlusion. During occlusion, precapillary sphincters exhibited pronounced Ca²⁺ elevations and constriction, amplifying downstream capillary resistance. Following reperfusion, elevated Ca²⁺ signals persisted without proportional diameter changes, indicating early uncoupling between Ca²⁺ dynamics and vascular responses. In the chronic phase, loss of precapillary sphincters-associated contractile pericytes was associated with capillary dilation and persistent neurovascular uncoupling. Although partial recovery of pericyte coverage and Ca²⁺ activity occurred, stimulus-evoked vascular responses remained blunted. These findings highlight precapillary sphincters as a key contributor to ischemia-induced microvascular dysfunction.

## Introduction

Ischemic stroke, characterized by occlusion of a cerebral artery causing insufficient blood flow and tissue infarction, remains a leading cause of global disability and mortality.^1^ However, even with successful recanalization of the occluded vessels, persistent microvascular dysfunction, such as neurovascular uncoupling^2–4^ and the ‘no reflow’^5–8^ phenomenon, continues to limit effective reperfusion and exacerbate brain injury, contributing to expansion of the ischemic core.

Neurovascular coupling (NVC) refers to the regional regulation of cerebral blood flow in response to rises in neuronal activity. Vascular mural cells are the contractile element to regulate vessel diameters and blood flow, which include vascular smooth muscle cells (VSMCs) in arteries and arterioles, as well as precapillary sphincters (PSs) and contractile pericytes in capillaries and venules.^9,10^ VSMCs , PSs, and contractile pericytes at the arteriole-capillary transitional (ACT) zone play a major role in blood flow regulation during NVC.^11–13^ Furthermore, the conventional opinion that pericytes in the capillary bed, venules, and veins are non-contractile has been challenged by recent studies, showing the presence of ɑ-smooth muscle actin (ɑ-SMA)^14^, which provide a mechanism for a tonic regulation of the vasculature^15,16^.

In ischemic stroke *in vivo* and during oxygen–glucose deprivation *in vitro,* pericytes undergo contraction and ‘die-in-rigor’ during both occlusion and reperfusion.^12^ In the chronic phase, prolonged contraction by pericytes at capillaries promotes ‘no flow’ phenomena and exacerbates red blood cell stalling.^17^ However, these studies have largely considered pericyte behavior across the broad capillary bed, and it remains unclear how ischemic stroke affects contractile pericytes in the ACT zone of the ischemic penumbra over days to weeks after the stroke.

PSs are located at the transition zone between the penetrating arterioles (PAs) and the first-order capillaries. They are characterized by thick, strongly contractile vascular mural cells that enwrap an indented vessel lumen with inner diameter of 3–4 µm.^18^ Positioned at the ACT zone, PSs act as gatekeepers of downstream capillary blood flow, protecting downstream capillaries from high pressure, and exerting a disproportionately large influence on the magnitude of local blood flow fluctuations.

Although PSs share many vasoactive signaling pathways with VSMCs and pericytes, they demonstrate superior responsiveness compared to VSMCs and contractile pericytes during NVC. PSs contract and dilate in response to the application of vasoactive compounds and are capable of both strong dilation (30–60%) as well as constriction (∼80%).^13,18,19^ Notably, PSs exhibit heightened vulnerability to pathological events like cortical spreading depolarization and aging, exhibiting greater increases in flow resistance and desensitization of signaling by local puff of vasoactive compounds.^18,20^ Despite their obvious importance, the role of PSs and their associated contractile pericytes in microvascular dysfunction induced by ischemic stroke has not been systematically investigated.

Mural cells regulate vascular tone and contribute to rapid NVC responses, primarily via intracellular calcium (Ca²⁺) signaling and actin–myosin light-chain–dependent contractile mechanisms.^21,22^ However, a comprehensive examination of functional Ca²⁺ dynamics of VSMCs, PSs, and contractile pericytes during stroke progression, in both acute and chronic phases, is lacking. Measurements of mural cell Ca²⁺ dynamics and vascular responsiveness in awake, behaving mice are more physiologically relevant than those obtained under anesthesia. In the current study we performed such measurements over 4 weeks, i.e. we characterized Ca²⁺ signaling and contractility of VSMCs, PSs, and contractile pericytes during both the acute and chronic phases of ischemic stroke.

Our results reveal that PSs play a pivotal role in the pathophysiology of ischemic stroke. During the acute phase, PSs and adjacent pericytes exhibit elevated levels of Ca²⁺, leading to excessive capillary constriction. This is followed by chronic pericyte loss, capillary dilation and impaired vascular reactivity. In contrast, contractile pericytes not associated with PSs demonstrate greater survival and better preservation of vascular tone and NVC responses. Although pericyte coverage and Ca²⁺ signaling partially recovered at later stages, activity-evoked vascular responses and Ca²⁺ sensitivity remained impaired, resulting in spatially heterogeneous NVC responses.

## Results

### Excessive Calcium Rise in Mural Cells, Especially PSs, Induces Strong Capillary Constriction During the Acute Phase of Ischemic Stroke

To investigate the effects of PSs during the acute phase of ischemic stroke, we performed craniotomy and transient middle cerebral artery occlusion (MCAO) model on Acta2-GCaMP8 mice, expressing genetically encoded green, fluorescent Ca^2+^ indicators in contractile vascular mural cells (**Fig. 1**). We recorded region of interests (ROIs) in the ACT zone revealing vascular mural cell Ca^2+^ dynamics together with vessel diameter changes in layer II/III of the somatosensory cortex, a region selected to sample peri-infarct cortex based on anatomical location and 2,3,5-triphenyltetrazolium chloride (TTC) reference staining (**Supplemental Fig. 1A**). As described in methods, PSs were identified based on their anatomical location at the arteriolar–capillary junction, characteristic focal indentation in vessel diameter, and morphology of surrounding mural cells^18^. We mapped PS locations within the cranial window, and imaged ACT zones with and without PSs at baseline, during occlusion (5 min after insertion of filament), and 15 min after reperfusion. Further, we classified capillaries by their branching orders, with the first order being the first capillary branching from the PA, and so forth (**Fig. 1A, B, Supplemental Fig. 2A**).

**Figure 1.**
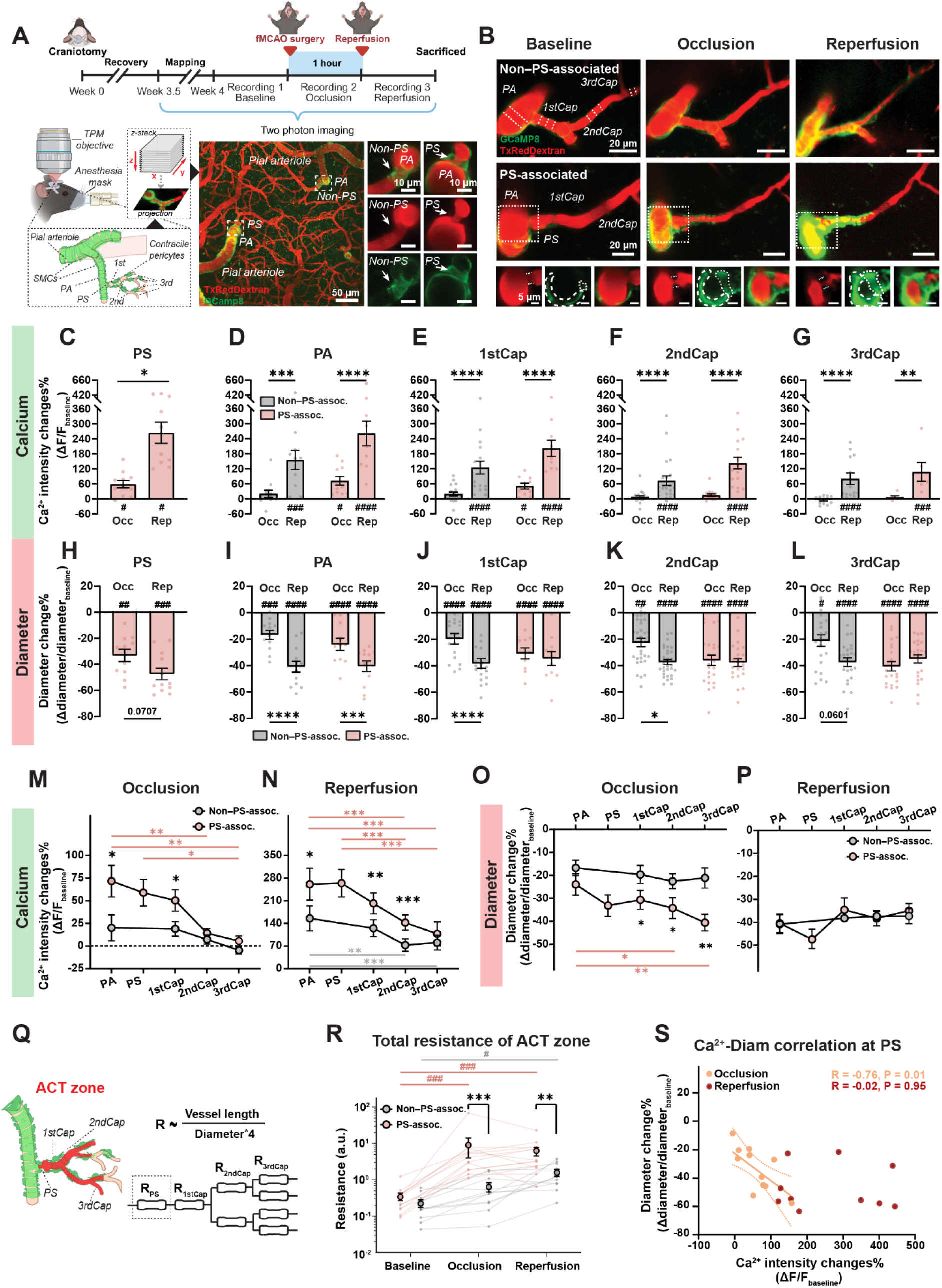
Ischemic stroke induces robust calcium elevation in mural cells and pronounced vascular constriction. (A) Experimental timeline of acute phase recording (top), schematic of the two-photon imaging strategy (bottom left), and representative images illustrating ROIs with precapillary sphincters (PS; PS-associated group) and without PSs (non−PS-associated group) (bottom right; scale bar, 50 µm). Penetrating arterioles (PAs), capillaries, and mural cells were acquired as z-stacks using two-photon imaging. (B) Representative imaging of non−PS-associated (top row) and PS-associated groups (middle row) at baseline, occlusion, and following reperfusion (scale bar: 20 μm). PSs are shown in the magnified imaging (bottom row, scale bar: 5 μm). Red: Texas Red–dextran (vessels); green: GCaMP8 fluorescence in Acta2-positive mural cells. (C–L) Ca^2+^ intensity (C–G) and diameter (H–L) changes in PSs, PAs, and first- to third-order capillaries, at different time points (n = 6–30 ROIs from 5 mice). Pink: PS-associated group; Grey: non−PS-associated group. (M–P) Comparison of Ca^2+^ signals (M, N) and diameter changes (O, P) between PS-associated (pink) and non−PS-associated (grey) groups during occlusion and reperfusion across different vascular segments (n = 6–30 ROIs from 5 mice). (Q, R) Schematic illustrating the calculation of vessel resistance in the arteriole-capillary transitional (ACT) zone (Q) and total resistance of the ACT zone (R) in PS-associated (pink) and non−PS-associated (grey) groups across different time points (n = 12 ROIs from 5 mice). (S) Scatter plot showing the relationship between Ca^2+^ and diameter changes during occlusion and reperfusion in PSs (n = 10–11 ROIs from 5 mice). Light orange: occlusion; dark orange: reperfusion. Data are presented as mean ± SEM. Statistical analyses included repeated-measures one-way ANOVA for panels C and H, linear mixed-effects model for panels D–G, I–L, M–P, and R. Pearson correlation analysis was used for panels S. Comparisons between baseline and post-stroke time points are indicated by # *P* < 0.05, ## *P* < 0.01, ### *P* < 0.001, and #### *P* < 0.0001. All other comparisons are indicated by * *P* < 0.05, ** *P* < 0.01, *** *P* < 0.001, and **** *P* < 0.0001.

Firstly, we compared Ca^2+^ and vascular dynamics during occlusion versus reperfusion. During occlusion, Ca^2+^ intensity in PSs increased by 58.9 ± 14.6%, which caused a 33.2 ± 4.7% reduction in vessel diameter. After reperfusion, Ca^2+^ intensity in PSs increased to 264.5 ± 42.3%, accompanied by a 47.4 ± 4.5% vasoconstriction (**Fig. 1C, H**). Similar trends were observed in PAs and first- to third-order capillaries within the ACT zone: compared with occlusion, reperfusion induced a markedly larger rise of mural cell Ca^2+^ in regions both with (PS-associated group) or without PSs (non–PS-associated group) (**Fig. 1D–G**). However, the vasoconstriction during occlusion and reperfusion differed between ACT zones in the PS-associated and non–PS-associated groups. In the non–PS-associated group, occlusion induced moderate vasoconstriction, which became more pronounced immediately after reperfusion. In contrast, in the PS-associated group, occlusion caused a strong reduction in capillary diameter in first- to third-order capillaries, which was comparable in magnitude to that observed during reperfusion (**Fig. 1I–L**). These results indicated that PSs were associated with an early, pronounced Ca^2+^ elevation and larger contraction in adjacent mural cells, which led to microvascular impairment in the acute phase of transient ischemic stroke.

Next, we examined whether there were differences in the Ca²⁺ rise and vasoconstriction between PS-associated and non-PS-associated segments. During occlusion, PS-associated segments showed greater mural cell Ca^2+^ elevations than non–PS-associated segments in both PAs and first-order capillaries. In the PS-associated segments, mural cells in PAs and PSs exhibited larger Ca²⁺ increases than pericytes in capillaries, a pattern not observed in non–PS-associated segments (**Fig. 1M**). During reperfusion, Ca^2+^ elevations remained greater in PS-associated segments, with more pronounced increases observed in PAs and PSs (**Fig. 1N**). In contrast, vasoconstriction patterns were different: during occlusion, vasoconstriction was more pronounced in first- to third-order capillaries in PS-associated segments. Furthermore, within PS-associated segments, vasoconstriction was stronger in second- and third-order capillaries than in PAs, a pattern not seen in non–PS-associated segments (**Fig. 1O**). During reperfusion, PS-associated and non–PS-associated segments exhibited similar degrees of constriction (**Fig. 1P**). Together, these results suggested that PSs amplified downstream capillary constriction.

To further assess whether ACT zones differ in resistance to blood flow depending on the presence of PSs, we adapted Poiseuille’s law (**Supplemental Fig. 3A**) to calculate total ACT vascular resistance using the series and parallel circuit configuration shown in **Fig. 1Q**. Although ACT zones in PS and non–PS-associated groups had similar baseline resistance (**Fig. 1R**; **Supplemental Fig. 3B**), occlusion–reperfusion caused an earlier and greater increase in resistance in ACT zones containing PSs. Furthermore, ACT zones in the PS-associated group showed higher resistance during both occlusion and reperfusion than those in the non–PS-associated group (**Fig. 1R**). This result suggested that heterogeneity in ACT zone resistance during the acute phase of ischemic stroke related to the presence or absence of PSs.

To quantify the relationship between the magnitudes of Ca^2+^ elevation and vasoconstriction, we performed Pearson correlation analysis at different time points. During occlusion, Ca^2+^ and diameter changes in PSs exhibited a strong negative correlation, consistent with Ca^2+^-dependent constriction; however, this relationship was disrupted following reperfusion (**Fig. 1S**). The pronounced Ca^2+^ elevation during reperfusion without additional constriction suggested Ca^2+^ dysregulation and saturated contraction in mural cells. Together, these findings suggested an early disruption of Ca^2+^-dependent vascular control in PS-containing ACT zones during the acute phase of ischemic stroke.

### PS-Associated Loss of Contractile Pericyte Coverage Post Ischemic Stroke

An abnormal increase in intracellular Ca²⁺ could be cytotoxic, activating calpains that degrade cytoskeletal and organelle proteins, and causing mitochondrial dysfunction through Ca²⁺ accumulation, loss of membrane potential and caspase activation, potentially leading to cell death and structural remodeling after ischemia.^23–25^ Previous studies have demonstrated that while the initial wave of pericyte death occurs acutely, the maximum reduction in pericyte coverage and the peak of delayed cell death (including necroptosis) in the capillary bed of peri-infarct regions are most pronounced between days 3 and 7.^18,26,27^ After day 7, robust angiogenesis^26,28^ and pericyte proliferation^27,29,30^ are typically observed. To investigate PS-associated microvascular impairments after stroke, we performed immunohistochemical staining on mouse brains collected at 4 days post-stroke. Vascular mural cells were labeled by PDGFR-β, contractile pericytes, VSMCs and PSs were labeled using α-SMA, while vascular endothelial cells were labeled by podocaylxin (**Fig. 2A**).

**Figure 2.**
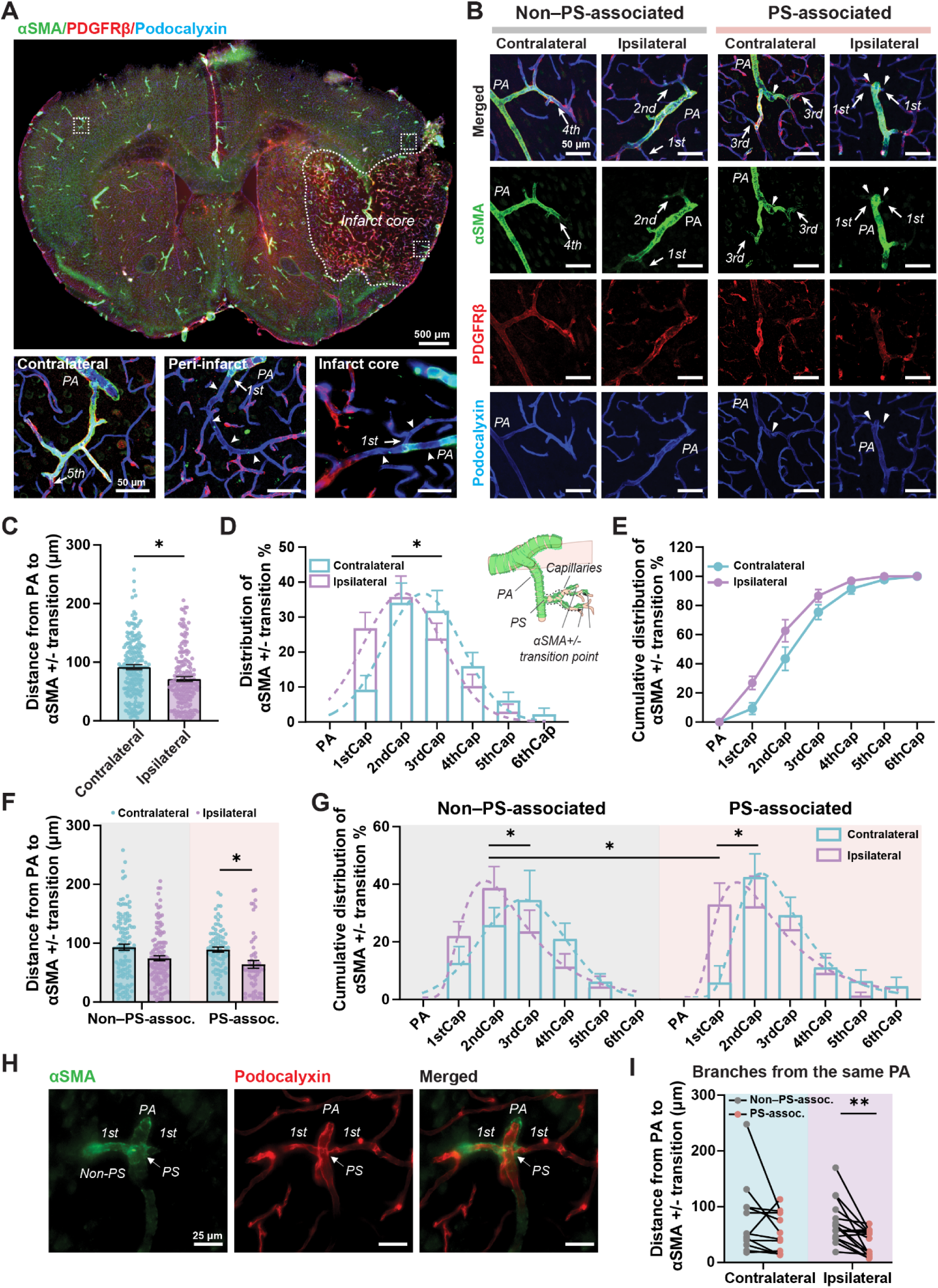
Ischemia-induced reduction of contractile pericyte coverage at day 4 post-stroke, analyzed by immunohistochemistry. (A) Overview of the coronal brain section showing the contralateral (left) and ipsilateral (right) hemispheres (scale bar: 500 µm), labeled with α-SMA (green), PDGFR-β (red), and podocalyxin (blue). Magnified images show representative regions from the contralateral (left), peri-infarct (middle), and ischemic core (right) areas (scale bar: 50 µm). Arrows indicate the endpoints of contractile pericyte coverages, and arrowheads indicate regions exhibiting loss of contractile pericyte coverage. (B) Representative images of vessels with precapillary sphincters (PSs, PS-associated) and without PSs (non−PS-associated) in contralateral and ipsilateral hemispheres (scale bar: 50 µm). Arrows denote the endpoints of contractile pericyte coverage, and arrowheads indicate the location of PSs. PA: penetrating arteriole; 1st−4th: first- to fourth- order capillaries. (C) Quantification of the distance from the junction of PA and first-order capillary to the endpoint of contractile pericyte coverage, comparing vessel segments from contralateral (blue) and ipsilateral (purple) sides (n = 184–202 ROIs from 5 mice). (D, E) Distribution (D) and cumulative distribution (E) of branch order at the endpoint of contractile pericyte coverage, comparing vessel segments from contralateral (blue) and ipsilateral (purple) sides. (F, G) Comparison of coverage distance (F) and capillary order at the endpoint (G) between PS-associated and non−PS-associated groups (n = 57–127 ROIs from 5 mice). Blue and purple indicate contralateral and ipsilateral sides, respectively. (H, I) Representative images of vessel pairs with and without PSs arising from the same PA (H), along with quantitative comparison (I) of coverage distances in contralateral and ipsilateral hemispheres (n = 12–16 vessel pairs from 5 mice). Pink and grey indicate PS-associated and non−PS-associated groups, respectively. Scale bar: 25 µm. Data are presented as mean ± SEM. Statistical analyses included a paired t-test for panels C, Bayesian analysis for panels D and G, and linear mixed-effects model for panels F and I. Statistical significance in panels C, F, and I: * *P* < 0.05, ** *P* < 0.01. For panels D and G, * denotes a statistically supported conclusion with posterior probability >0.95.

To quantify contractile pericyte coverage, we measured the distance from PA–first-order capillary junction to the distal endpoint of contractile pericyte coverage (α-SMA–positive coverage), as well as the capillary order at the endpoint. In general, contractile pericyte coverage distance was significantly shorter on the ipsilateral side (stroke) compared to the contralateral side (**Fig. 2B–C**). Accordingly, the capillary order at the endpoint of contractile pericyte coverage was lower on the ipsilateral side (**Fig. 2D–E**), reflecting retraction and loss of contractile pericyte coverage in the ACT zone.

When further stratified by the presence of PSs, reductions in coverage distance and terminal capillary order of contractile pericytes were significantly more pronounced in PS-associated group as compared with non−PS-associated segments, which exhibited a moderate decrease in coverage distance (**Fig. 2F–G**). Interestingly, when comparing branches in the PS-associated and non−PS-associated segments arising from the same PA, coverage distance was also significantly shorter in PS-associated segments on the ipsilateral side, but not on the contralateral side post-stroke (**Fig. 2H, I**). These results suggested that ischemic stroke reduced contractile pericyte coverage, and this effect was intensified in vascular segments with a PS.

### Longitudinal *In Vivo* Imaging Reveals Progressive Pericyte Coverage Loss and Capillary Responsivity

Given the marked Ca^2+^ surge in PS-associated regions during the acute phase and the reduction in contractile pericyte coverage revealed by immunohistochemistry in the first week post-stroke, we hypothesized that PSs were associated with a greater likelihood of contractile pericyte pathology and dysfunction after stroke. To test this hypothesis, we performed longitudinal two-photon microscopy (TPM) and laser speckle contrast imaging (LSCI) in awake Acta2-GCaMP8 mice. Fast repetitive volumetric (x–y–z–t) hyperstack imaging was used to capture the full vascular architecture and dynamic mural cell activity over time. We recorded Ca^2+^ signals of VSMCs at PAs, PSs and contractile pericytes, as well as their corresponding vascular diameters during control conditions and in response to whisker air puff stimulation (**Fig. 3A, B**).

**Figure 3.**
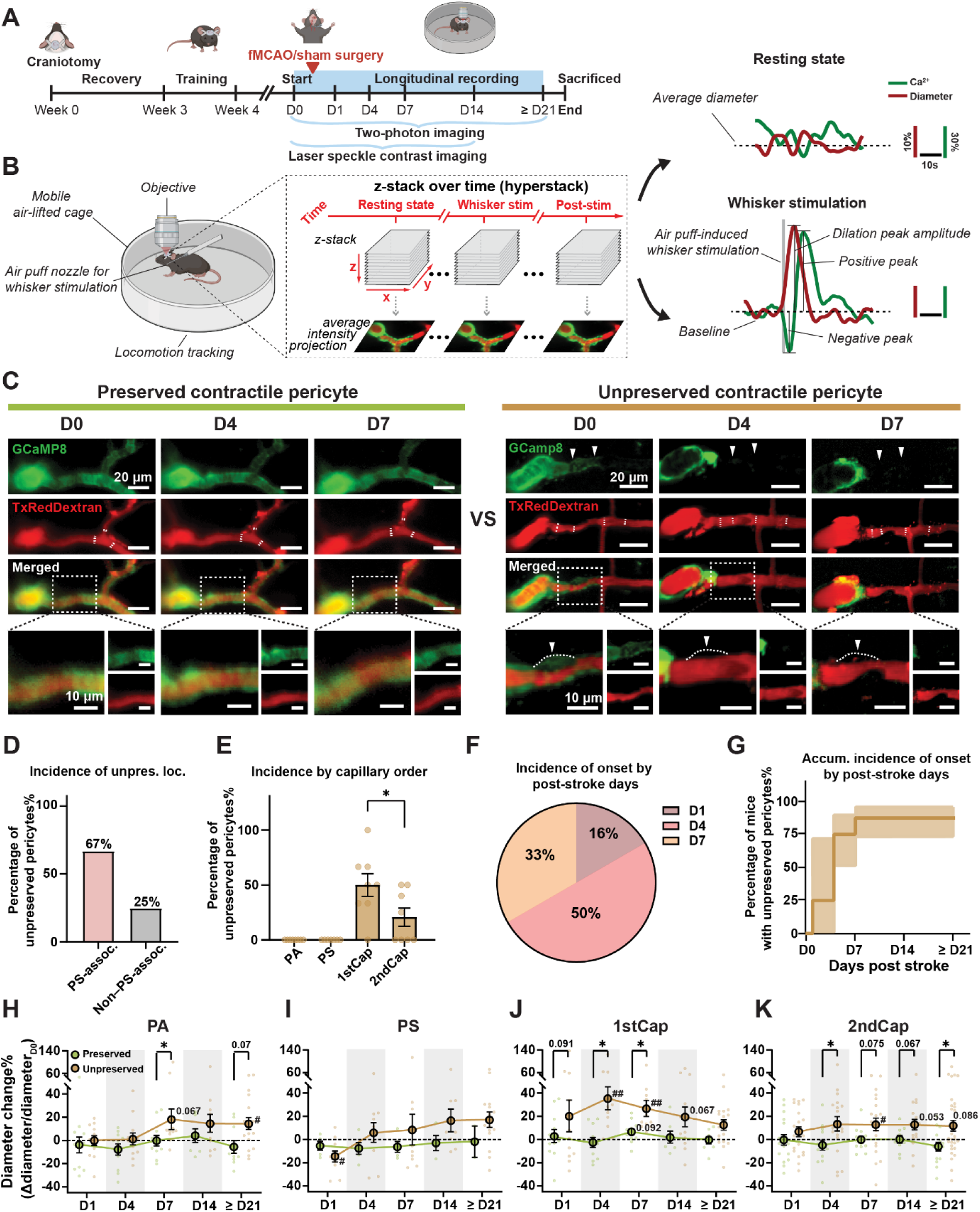
Long-term *in vivo* imaging reveals loss of pericyte calcium signaling accompanied by progressive capillary dilation following ischemic stroke. (A) Experimental timeline for chronic-phase awake imaging in Acta2-GCaMP8 mice after ischemic stroke. fMCAO, filament middle cerebral artery occlusion. (B) Schematic of awake two-photon imaging (left), imaging strategy (middle), and representative changes in vascular diameter and Ca^2+^ intensity during resting state and whisker responses (right). Fluctuations during the resting state and responses evoked by whisker stimulation in vessels and mural cells were recorded using an x-y-z-t hyperstack. Z-projected signals across time were then quantified to generate diameter and Ca^2+^ response curves during resting and stimulation periods. (C) Representative images of ROIs showing preserved and unpreserved contractile pericyte Ca^2+^ signals at various days post-stroke (Scale bar: 20 µm). The red channel shows Texas Red-dextran-labeled vasculature, and the green channel shows GCaMP8 fluorescence. The magnified image highlights first-order capillaries (Scale bar: 10 µm). (D) Percentage of vessels with unpreserved contractile pericyte in region with precapillary sphincters (PSs, PS-associated; pink) and without PSs (non−PS-associated; grey). (E) Percentage of unpreserved contractile pericytes across different vessel segments (N = 8 mice). (F, G) Incidence of unpreserved pericytes (F) and percentage of mice exhibiting unpreserved pericytes (G) at different post-stroke time points (N = 8 mice). (H–K) Comparison of percentage changes in vessel diameter over time between preserved (green) and unpreserved (brown) groups across penetrating arterioles (PAs) (H), PSs (I), first-order capillaries (1stCap) (J), and second-order capillaries (2ndCap) (K). Data represent n = 7–18 (PAs), 2–15 (PSs), 7–18 (first-order capillaries), and 10–31 (second-order capillaries) ROIs from 8 mice. Data are presented as mean ± SEM. Panel D was shown descriptively. A linear mixed-effects model was applied for panel E, and H–K. Comparisons between D0 and post-stroke time points are indicated by # *P* < 0.05, ## *P* < 0.01. All other comparisons are indicated by * *P* < 0.05.

Starting as early as one day post-stroke, we observed a progressive loss of Ca^2+^ signals, most prominently in contractile pericytes on first-order capillaries (**Fig. 3C**). To classify pericyte Ca^2+^ signal loss, we quantified Ca^2+^ coverage relative to baseline (D0), as well as foreground-background contrast. Pericytes were classified as unpreserved based on a sustained reduction in Ca²⁺ signal–defined coverage (<50% of baseline), loss of foreground signal to background levels, preserved signal-to-noise, and concurrent local vessel dilation (>20% from baseline), ensuring that coverage loss reflected true local signal loss rather than imaging artifacts. ROIs not fulfilling all criteria were classified as preserved pericytes (**Supplemental Fig. 4**). Detailed criteria were described in the Methods. We then calculated the proportion of unpreserved pericytes in both PS-associated and non−PS-associated segments. Descriptively, first-order capillaries in PS-associated segments more frequently exhibited unpreserved pericytes than those in non−PS-associated segments (67% versus 25%; **Fig. 3D**). In sham mice, pericyte coverage appeared stable across imaging sessions, arguing against a major effect of longitudinal imaging alone (**Supplemental Fig. 5A-H**).

Quantification across vessel segments and time points revealed that unpreserved pericyte occurred predominantly in first-order capillaries during the first week following ischemia (**Fig. 3E–G**). We also observed unpreserved pericytes associated with second-order capillaries; however, the proportion of unpreserved pericytes in second-order capillaries was significantly lower than that in first-order capillaries (**Fig. 3E**). Subsequent analyses focused on the unpreserved pericytes located in first-order capillaries.

Unpreserved contractile pericytes may affect the vessel diameter in the resting state. To assess this hypothesis, we measured resting-state vessel diameters across different vascular segments over time. Measurements were stratified according to the preservation status of pericytes in the first-order capillaries. Capillaries and PAs in the unpreserved group exhibited significant dilation post stroke, whereas resting diameters remained unchanged in the preserved group (**Fig. 3H, J, K**). These findings are consistent with previous studies showing loss of basal tone in regions with laser-ablated or age-related coverage loss of pericytes.^31,32^ Although a trend toward vasodilation was also observed at PSs in the unpreserved group, these changes did not reach statistical significance (**Fig. 3I**). These findings suggested that an ischemic stroke induced PS-associated impairment of contractile pericytes in the chronic phase, contributing to impaired regulation of vascular tone and capillary dilation.

### Calcium and Vascular Responses Differ Between Unpreserved and Preserved Pericytes Under Healthy Conditions

Having demonstrated reduced contractile pericyte coverage and increased resting-state vessel diameter during the chronic post-stroke phase, we next examined whether pericytes that later became unpreserved exhibit distinct NVC responses at baseline (D0, healthy state) compared with those that remained preserved after ischemic stroke. For this purpose, we retrospectively stratified pericytes at D0 based on their post-stroke fate and compared mural cell Ca^2+^ responses and vascular diameter changes evoked by air-puff–induced whisker stimulation in awake Acta2-GCaMP8 mice (**Fig. 3A, B**).

At D0, i.e. prior to the stroke, all vascular segments in both preserved and unpreserved groups exhibited robust Ca^2+^ responses that effectively drove vessel dilation in response to whisker stimulation (**Fig. 4**). However, pericytes in first- to second-order capillaries showed reduced Ca^2+^ responses following whisker stimulation in the unpreserved group compared with preserved group (**Fig. 4A−C**). Consistent with these findings, whisker stimulation–induced vascular dilation was attenuated in the unpreserved group relative to the preserved group (**Fig. 4D, E**). In contrast, baseline vessel diameters did not differ between groups across vascular segments (**Fig. 4F).**

**Figure 4.**
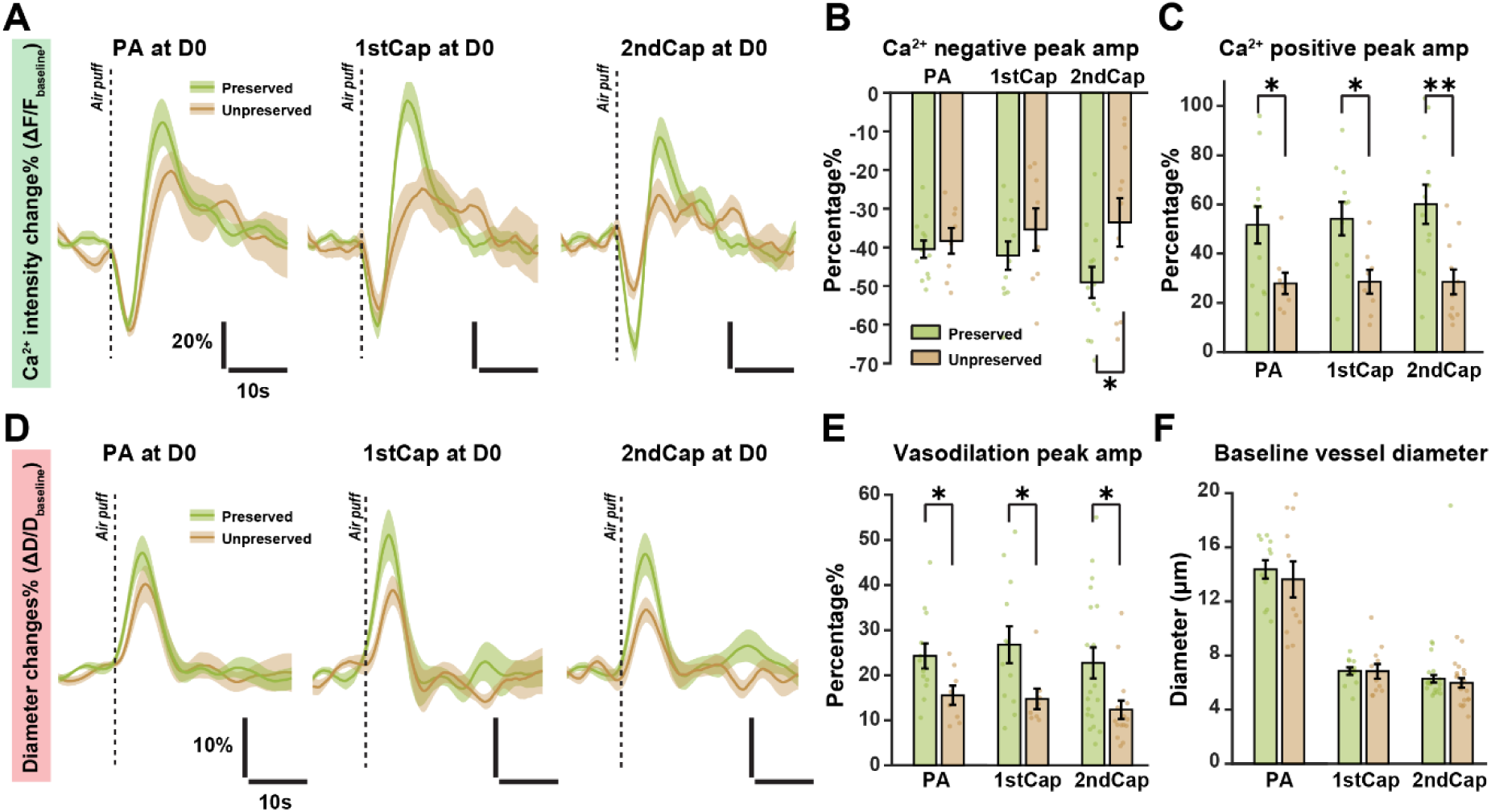
Unpreserved contractile pericytes and their corresponding vessel segments exhibit distinct calcium and vascular responses to whisker stimulation at D0. (A, D) Ca^2+^ (A) and vascular (D) response curves in different mural cells and corresponding vessel segments in the preserved (green) and unpreserved (brown) groups at D0. Each curve represents the mean response averaged across eight mice. (B, C) Amplitude of negative (B) and positive (C) Ca^2+^ peaks in penetrating arterioles (PAs), first-order capillaries (1stCap) and second-order capillaries (2ndCap) at D0 in the preserved (green) and unpreserved (brown) groups, normalized to the pre-stimulation baseline (25 s before air-puff induced whisker stimulation). Data represent n = 18−24 ROIs from 8 mice. (E) Amplitude of peak vascular responses to whisker stimulation in PAs, first-and second-order capillaries at D0, normalized to the pre-stimulation baseline (25 s before air-puff induced whisker stimulation). Data represent n = 19−34 ROIs from 8 mice. (F) Baseline vessel diameter in PAs, first-and second-order capillaries at D0. Data represent n = 22−44 ROIs from 8 mice. Data are presented as mean ± SEM. A linear mixed-effects model was applied for panels B, C, E, and F. * *P* < 0.05, ** *P* < 0.01.

Interestingly, when the data were stratified based on the presence of PSs, reduced Ca^2+^ responses were also observed in pericytes of first- to second-order capillaries within PS-associated segments compared with non−PS-associated segments (**Supplemental Fig. 6A−C**). In addition, vascular dilation in response to whisker stimulation was smaller in PS-associated segments than in non−PS-associated segments (**Supplemental Fig. 6D, E**), whereas baseline vessel diameters showed no difference between groups (**Supplemental Fig. 6F**). These findings indicated that PS-associated ACT regions exhibited attenuated Ca²⁺ and vasodilatory responses at baseline despite the known strong contractile capacity of PSs. These findings were consistent with the differences observed between preserved and unpreserved groups.

In summary, these results reveal a similar pattern of whisker-evoked responses in mural cells and vessel segments between PS-associated and unpreserved groups, both of which differ markedly from non−PS-associated and preserved groups, respectively. These findings may suggest that mural cells within PS-associated ACT regions exhibit distinct NVC responses in healthy state, potentially predisposing them to pericyte dysfunction following ischemic stroke and during the chronic phase.

### Attenuated Calcium Dynamics Impair NVC and Vessel Diameter Changes at Contractile Pericytes

After characterizing the differences between preserved and unpreserved groups before ischemic stroke, we next examined whether their capacity to regulate blood flow differed after stroke in the chronic phase. Although most pericyte Ca²⁺ signals were undetectable in the unpreserved group, Ca²⁺ could still be quantified based on the residual fragments of pericytes in ROIs. When coverage decreased to <10% of baseline (D0), Ca²⁺ signals became too sparse for reliable quantification; therefore, these ROIs were excluded from analysis at that time point. Because this exclusion criterion preferentially removed the most severely affected ROIs, missingness at later time points was likely non-random and should be interpreted accordingly. Following ischemic stroke, both Ca^2+^ and vascular responses were markedly attenuated. Overall, Ca^2+^ responses were strongly suppressed during the first week post-stroke, with the exception that reductions in negative Ca^2+^ peak changes in PAs persisted up to day 21 (**Supplemental Fig. 7A–I**). In contrast to Ca^2+^ signaling, vascular responses to whisker stimulation across all vascular segments remained impaired for up to 21 days post-stroke, far outlasting the period of suppressed Ca^2+^ responses (**Supplemental Fig. 7J–N**).

To determine whether unpreserved pericytes contributed to these changes, we compared Ca^2+^ response amplitudes between preserved and unpreserved groups across post-stroke time points. Compared with D0, both groups exhibited pronounced reductions in Ca^2+^ response peaks in pericytes surrounding first-order capillaries during the first week (**Fig. 5A, B, E, F**). From the second week onward, Ca^2+^ responses in the remaining pericytes largely recovered. No significant differences in Ca^2+^ responses were observed between preserved and unpreserved groups across time points (**Fig. 5B, E, F**). In contrast, first-order capillaries in the unpreserved group exhibited a greater and more prolonged reduction in dilation (lasting until D21) compared with those in the preserved group (lasting until D14) (**Fig. 5C, D, G**).

**Figure 5.**
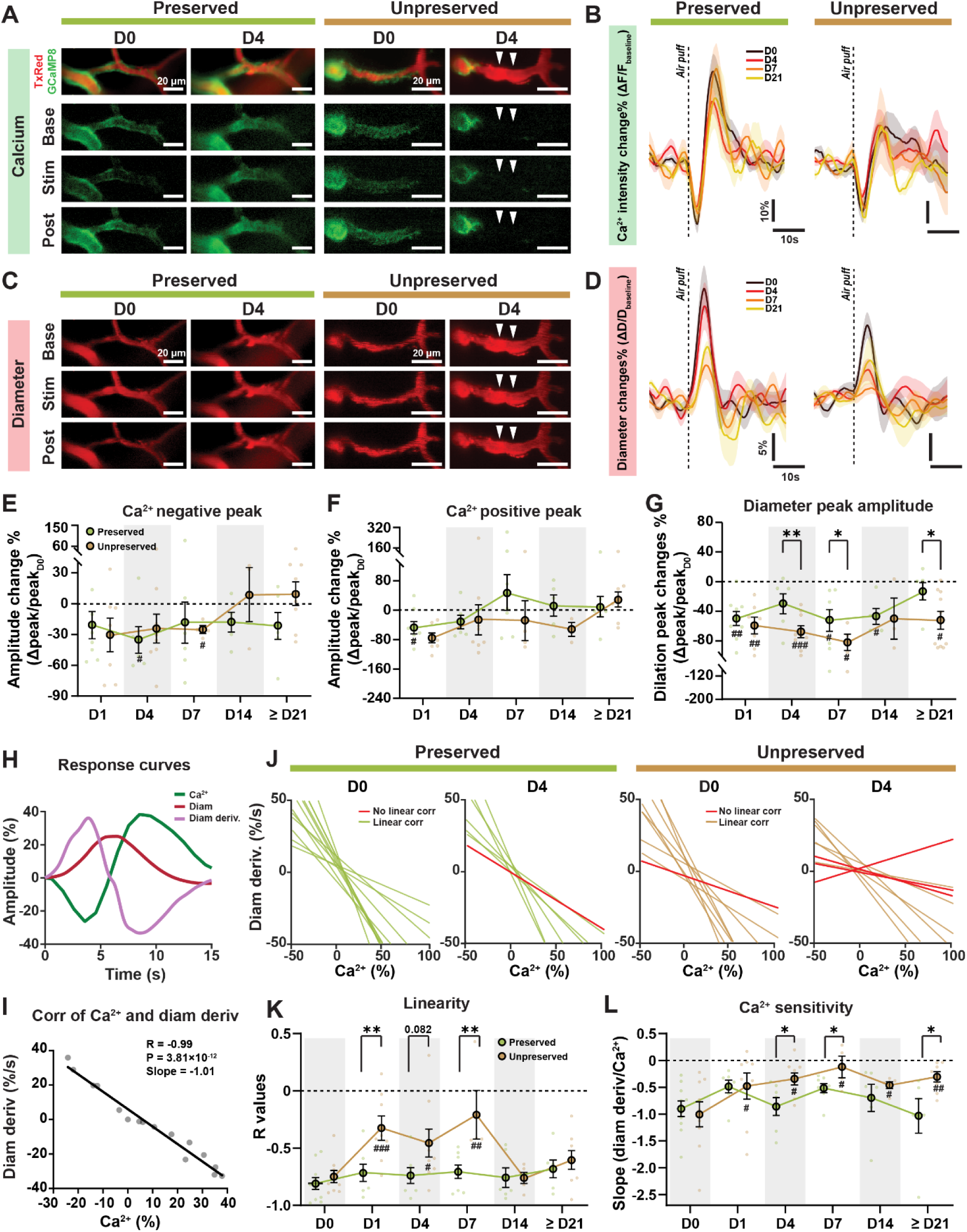
Ischemia reduces and uncouples mural cell calcium and vascular responses to whisker stimulation. (A, C) Representative images of whisker stimulation-induced mural cell Ca^2+^ (A) and vascular (C) responses in ROIs with preserved and unpreserved pericyte Ca^2+^ signals at D0 and D4 post-stroke (scale bar: 20 µm). Red: Texas Red–dextran; green: GCaMP8 fluorescence. Arrowheads indicate sites with absent Ca^2+^ signals. (B, D) Ca^2+^ and vascular response curves in pericytes and corresponding first-order capillaries from preserved and unpreserved groups (N = 8 mice), with post-stroke days color-coded. (E−G) Amplitude of negative Ca^2+^ peaks (E), positive Ca^2+^ peaks (F), and peak vascular responses (G) in first-order capillaries at different post-stroke time points, normalized to D0 (n = 3–8 ROIs for E–F; 3–9 ROIs for G; 8 mice). Green denotes the preserved group, and brown denotes the unpreserved group. (H) Representative traces of whisker stimulation-induced vascular diameter changes (red), mural cell Ca^2+^ responses (green), and the derivative of vascular diameter (purple) over time. (I) Scatter plots showing the relationship between Ca^2+^ signals and the time derivative of diameter changes. Pearson correlation coefficients (*R*), *P* values, and slopes of the linear regression lines are indicated. (J) Regression lines of Ca^2+^–vascular response correlations from 8 mice in preserved and unpreserved groups in first-order capillaries at D0 and ≥ D21. Green and brown lines indicate statistically significant correlations (*P* < 0.05), whereas red lines indicate non-significant correlations (*P* ≥ 0.05). (K, L) Summary of Pearson correlation coefficients (*R*; K) and slopes of regression lines (L) for correlations between Ca^2+^ signals and the derivative of vascular diameter in first-order capillaries at D0 and various days post stroke (n = 4−10 ROIs from 8 mice). Data are presented as mean ± SEM. Linear mixed-effects model was applied for panel E–G, K, L. Comparisons between D0 and post-stroke time points are indicated by # *P* < 0.05, ## *P* < 0.01, and ### *P* < 0.001. Comparisons between preserved and unpreserved groups are indicated by * *P* < 0.05, and ** *P* < 0.01.

To explore whether impairment of pericytes in first-order capillaries affects adjacent vessel segments, we classified whisker responses in PAs and second-order capillaries according to the preservation status of pericytes in first-order capillaries. In PAs, both Ca^2+^ and vascular responses were attenuated after stroke, with no significant differences between preserved and unpreserved groups (**Supplemental Fig. 8A, C, E**). In contrast, pericytes surrounding second-order capillaries in the unpreserved group exhibited a greater change in Ca^2+^ response peaks compared with the preserved group (**Supplemental Fig. 8B, D**). Second-order capillaries also showed prolonged and pronounced decrease in dilation amplitude (**Supplemental Fig. 8F**).

This suggests that sustained impairment of vascular responses cannot be explained by reduced Ca^2+^ responses in surviving mural cells but rather reflects coverage loss of functional contractile pericytes. Collectively, these results may demonstrate that ischemia leads to a disproportionate and prolonged impairment of NVC responses, particularly in regions with unpreserved contractile pericytes.

### Ischemia Caused Uncoupling Between Mural Cell Calcium Dynamics and Diameter Changes, Along with Reduction in Calcium Sensitivity

To further quantify the mismatch between Ca^2+^ and vascular responses during whisker stimulation, we performed Pearson correlation analysis to examine if ischemia disrupts the coupling between mural cell Ca^2+^ dynamics and vessel diameter changes. Elevations in Ca²⁺ activate the contractile machinery via calmodulin and myosin light chain kinase, leading to crossbridge cycling, active force generation, and vasoconstriction, whereas decreases in Ca²⁺ promote the opposite response.^22^ Therefore, we compared Ca^2+^ signals to the rate of diameter change (dD/dt), i.e., the dynamic mechanical response to Ca²⁺ transients. The derivative of diameter responses during whisker stimulation showed a strong inverse correlation with Ca^2+^ responses, whereas there was no or only a weak linear correlation between Ca^2+^ and diameter changes (**Fig. 5H, I**). This indicated that dD/dt captured the dynamic mechanical response to Ca²⁺ transients more sensitively than diameter alone.

We next examined this relationship across different vessel segments and time points after stroke. Ischemic stroke uncoupled Ca^2+^ signaling from vascular response, lasting from 1 day in PAs to up to 4–21 days in PSs and capillaries (**Supplemental Fig. 9A–E**). The slope of the regression line, which reflects the sensitivity of diameter changes to Ca^2+^ alterations, revealed a prolonged impairment in Ca^2+^ sensitivities within contractile pericytes up to D21 (**Supplemental Fig. 9A, H, I**). For PSs, reduction in Ca^2+^ sensitivity was only observed during the 1–4 days post stroke, whereas no significant change was detected in PAs (**Supplemental Fig. 9F, G**).

To determine whether unpreserved pericyte contributed to this uncoupling, we further stratified the data based on the preservation of pericyte Ca^2+^ signals at first-order capillaries. In first-order capillaries, both the linear correlation coefficient (*R* value) and the Ca^2+^ sensitivity (regression slope) were reduced to a greater extent and for a longer duration in the unpreserved group (**Fig. 5J–L**). In PAs and second-order capillaries, linearity was also impaired after ischemic stroke, particularly in the unpreserved group. Moreover, Ca^2+^ sensitivity in these vessel segments exhibited a greater reduction in the unpreserved group compared with the preserved group. (**Supplemental Fig. 10A–D**). These findings are consistent with our earlier observations (**Fig. 5A–G; Supplemental Fig. 8**), indicating that Ca^2+^ signaling in the survival mural cells is insufficient to effectively regulate vascular diameter once downstream contractile pericytes are unpreserved. These results suggested that an ischemic stroke induced a sustained uncoupling between mural cell Ca^2+^ dynamics and vascular responses, together with a reduction in Ca^2+^ sensitivity, particularly in capillary regions affected by unpreserved contractile pericytes.

### Pericyte Coverage Loss Led to Regional Ischemia During Functional Hyperemia

To directly assess how ischemia alters blood flow distribution at the mesoscopic level, we performed LSCI to measure cortical blood flow responses to whisker stimulation across a wider cortical field, i.e., the entire cranial window over the course of stroke progression in the same mice that had been examined with TPM. In the healthy state, approximately 84% of the imaged cortical area exhibited a robust and spatially widespread increase in blood flow following whisker stimulation (**Fig. 6A, B**). Following ischemic stroke, these responses were markedly attenuated, particularly during the first week post-stroke, as reflected by significant reductions in both response area and peak amplitude (**Fig. 6A–E**). Starting from day 14 post-stroke, blood flow responses partially recovered; however, the response amplitude remained significantly reduced compared with the healthy state (**Fig. 6D**).

**Figure 6.**
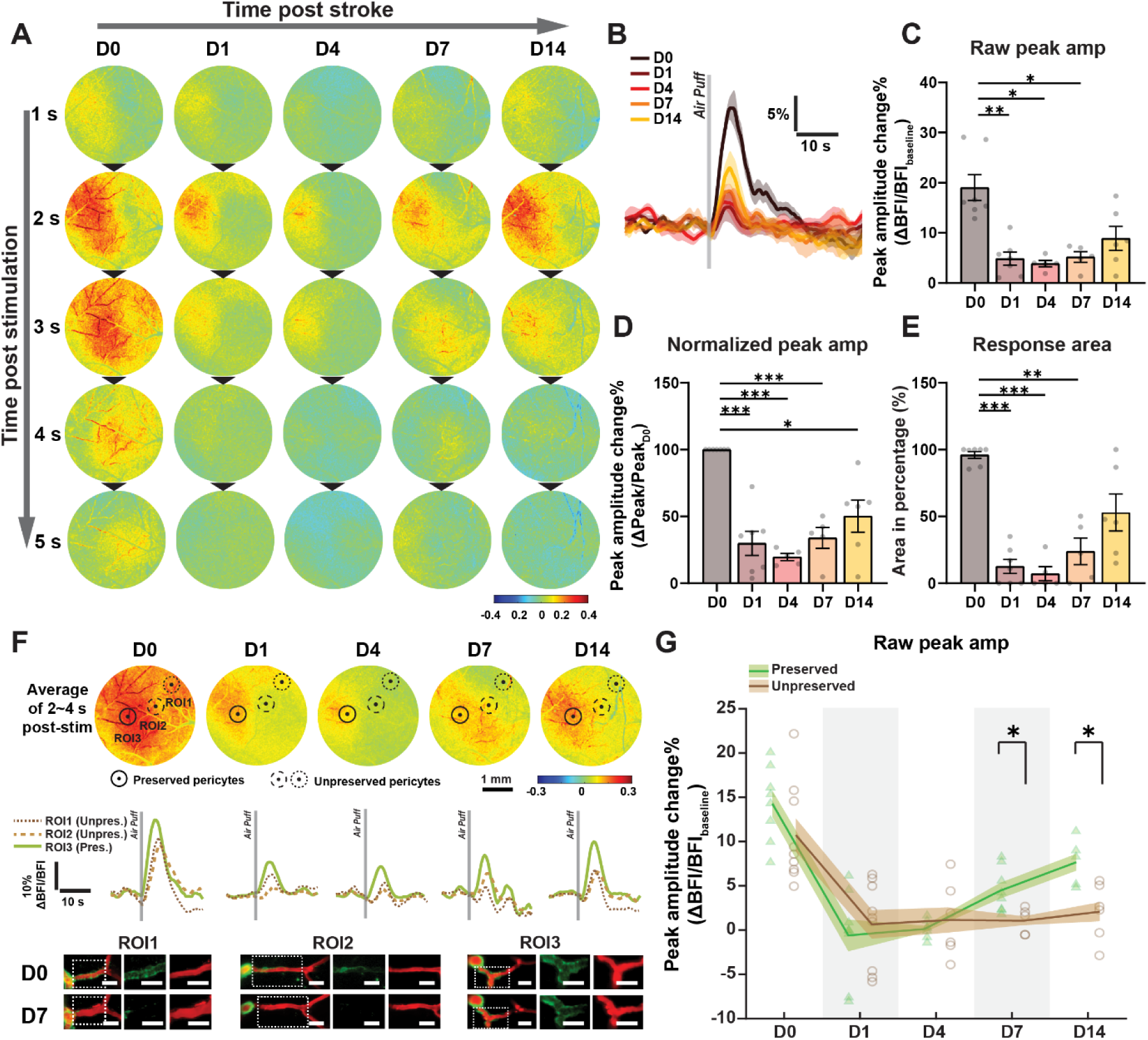
Regions associated with unpreserved pericytes exhibit a more severe impairment of whisker-evoked blood flow responses following ischemic stroke. (A) Representative heatmaps showing changes in blood flow index (BFI) measured 5 s after air-puff–induced whisker stimulation at D0, D1, D4, D7, and D14 post-stroke. BFI values were calculated as the difference between post-stimulation and pre-stimulation (baseline) signals. (B) Overlaid BFI response curves across different time points, with each day indicated by a distinct color. Each curve is the mean response overlaid from 6 mice. (C, D) Raw peak amplitudes (C) and normalized peak amplitudes (D) of BFI responses evoked by whisker stimulation (N = 7 mice). (E) Percentage of the responsive area relative to the total field of view at each time point (N = 6 mice). (F) Top row: Average heatmaps of BFI changes at baseline (D0), D1, D4, D7, and D14, calculated over 2–4 s after whisker stimulation (scale bar, 1 mm). Dashed circles (ROI1, ROI2) and hollow circles (ROI3) indicate unpreserved and preserved pericyte–associated regions, respectively. Middle row: BFI response traces from unpreserved pericyte–associated regions (brown dashed lines, ROI1 and ROI2) and preserved regions (green line, ROI3) during whisker stimulation across post-stroke time points. Bottom row: Representative TPM images showing pericyte coverage in ROI1–ROI3 at D0 and D7. (G) Comparison of raw peak BFI amplitudes evoked by whisker stimulation between preserved and unpreserved pericyte–associated regions across post-stroke time points (n = 5−9 ROIs from 7 mice). LSCI analyses were performed in a subset of the chronic imaging cohort (N = 7 of 8 mice). Data are presented as mean ± SEM. A linear mixed-effects model was applied for panels C−E, and G. * *P* < 0.05, ** *P* < 0.01, \*\*\**P* < 0.001.

To determine whether stroke-associated unpreserved pericytes at first-order capillaries contributed to the heterogeneous blood flow responses, we mapped ROIs previously monitored by TPM onto LSCI heatmaps using pial arteries as anatomical landmarks (**Fig. 6F**). Blood flow changes during whisker stimulation were quantified within circular regions (50 µm diameter) centered on TPM-recorded ROIs. The region size was selected based on the estimated distance of approximately 100 µm between two adjacent PAs in the mouse cortex.^33^ During the early post-stroke period (days 1–4), blood flow responses were similarly reduced in regions associated with preserved and unpreserved pericytes. However, from day 7 and onward, blood flow responses in regions associated with preserved pericytes showed partial recovery, whereas regions associated with unpreserved pericytes remained largely unresponsive to stimulation (**Fig. 6F, G**). These findings closely mirrored the TPM results and indicated that unpreserved pericytes disrupted local regulation of vessel diameter, leading to persistent impairment of blood flow regulation in adjacent cortical regions. In summary, the data suggested that following focal ischemia, NVC responses became spatially heterogeneous in the ischemic penumbra of awake mice, suggesting damage to contractile pericytes and failure of local neurovascular regulation.

### Recovery of Pericyte Coverage Does Not Fully Restore Whisker-Evoked Vascular Responses After Ischemic Stroke

Previous studies have shown that capillary pericytes can extend their processes to compensate for the loss of adjacent pericytes, leading to recovery of pericyte coverage within 21 days after ablation under physiological conditions.^31,32^ Consistent with this observation, we observed that ROIs exhibiting loss of pericyte Ca^2+^ signals during the first week after stroke showed partial restoration of pericyte coverage from the second week onward (**Fig. 7A**). To determine whether this structural restoration was accompanied by recovery of pericyte function, we classified each ROI into unpreserved and restored phases based on longitudinal changes in pericyte coverage at first-order capillaries. Onset of coverage restoration was defined as regrowth of pericyte coverage to either at least 20% above the minimum coverage observed after stroke or to more than 50% of baseline (D0) coverage. (**Supplemental Fig. 4D, H**). About 11.1% of ROIs didn’t show coverage restoration during the first month of TPM tracking, whereas nearly half of the restoration occurred at D21. The remaining ROIs started to restore at either 7 or 14 days post stroke (**Fig. 7B**). Furthermore, pericyte restoration in PS-associated and non−PS-associated groups was 70% and 50%, respectively (**Fig. 7C**).

**Figure 7.**
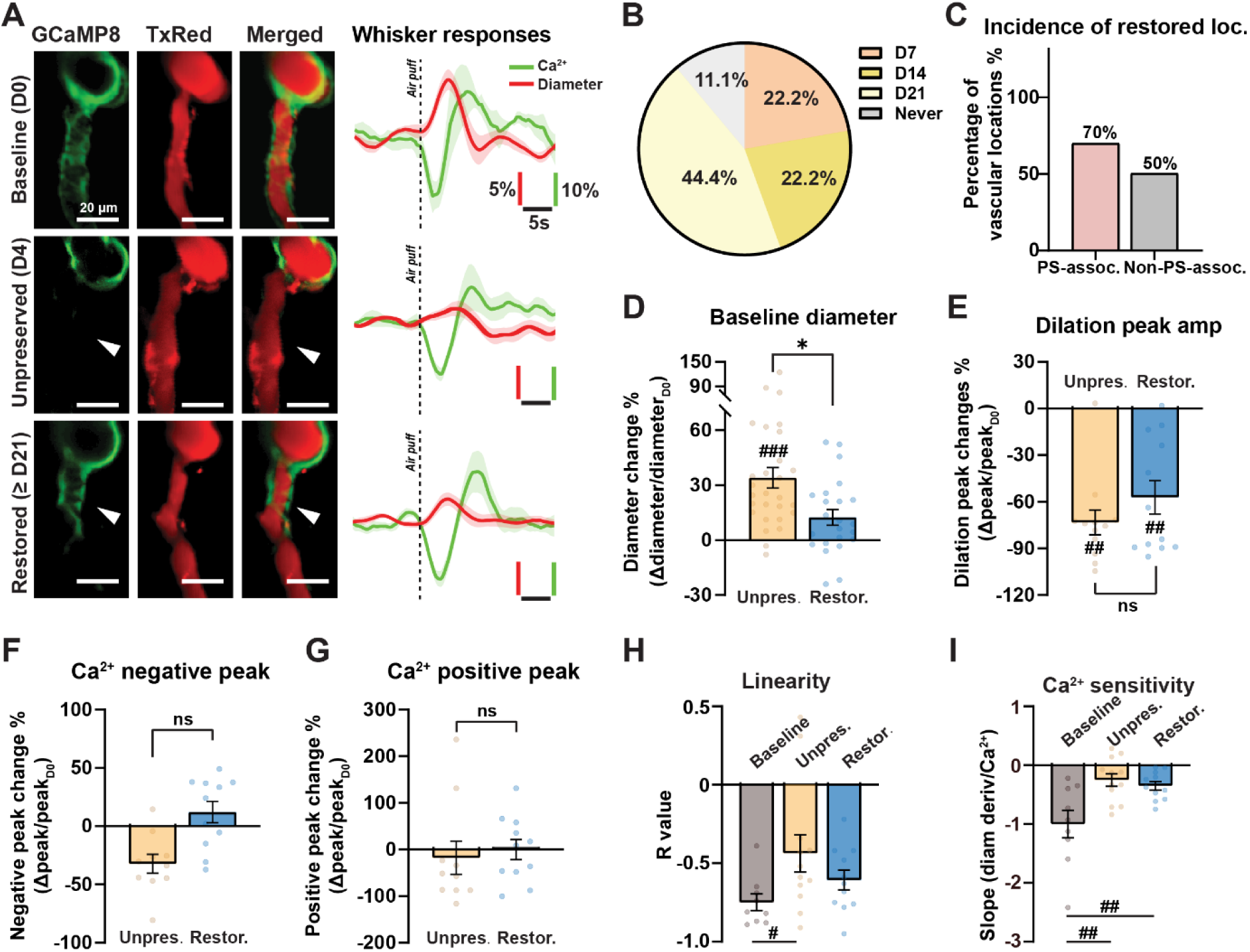
Recovery of pericyte coverage does not fully restore whisker-evoked vascular responses after ischemic stroke. (A) Representative image (left) and whisker responses (right) of ROIs showing pericyte coverage loss and subsequent restoration at baseline (D0), D4 (unpreserved pericyte phase), and ≥ D21 (pericyte restored phase). Red: Texas Red–dextran; green: GCaMP8 fluorescence. Arrowheads indicate sites of absent and restored Ca^2+^ signals. Scale bar: 20 µm. (B) The incidence of restoration at different days post stroke. Each day is indicated by a distinct color. (C) Percentage of restored ROIs in regions with precapillary sphincters (PSs; PS-associated) and without PSs (non−PS-associated). (D–G) Basal diameter changes (D), amplitude of peak vascular responses (E) and mural cell Ca^2+^ responses (F, G) in first-order capillaries during the unpreserved and restored phases (n = 10–28 ROIs from 8 mice). (H, I) Pearson correlation coefficients (*R*) between Ca^2+^ signals and the derivative of vascular diameter (H), as well as slopes of regression lines (I) in first-order capillaries during the baseline, unpreserved and restored phases (n = 8–13 ROIs from 8 mice). Data are presented as mean ± SEM. Panel B and C are shown descriptively. A linear mixed-effects model is applied for panels D–I. Comparisons between baseline and post-stroke time points are indicated by # *P* < 0.05, ## *P* < 0.01, and ### *P* < 0.001. Comparisons between unpreserved and restored phases are indicated by * *P* < 0.05.

During the time when pericyte function was affected, the basal diameters of capillaries increased, then returned toward baseline levels during the restoration phase (**Fig. 7D**). During the restoration phase, stimulation-induced mural cell Ca^2+^ responses were similar to responses at D0 (**Fig. 7F, G**). In contrast, whisker-evoked vascular dilation remained significantly impaired in first-order capillaries during both unpreserved and restored phases (**Fig. 7E**). Consistent with this observation, correlation analysis between the derivative of vessel diameter and Ca^2+^ signals revealed persistent impairment in the Ca^2+^ sensitivity but not correlation of first-order capillaries during the restored phase (**Fig. 7H, I**), indicating sustained dysfunction in vascular segments that previously experienced pericyte coverage loss. Together, these findings suggested that although pericyte coverage and Ca^2+^ signaling partially restored in the ischemic penumbra after ischemic stroke, this restoration was insufficient to fully restore NVC responses. Instead, ischemia induced a long-lasting reduction in sensitivity of vascular responses to mural cell Ca^2+^ during NVC responses, particularly in regions with unpreserved contractile pericytes.

## Discussion

In this study, we investigated the role of Ca²⁺-dependent contractility in mural cells, specifically SMCs, PSs, and contractile pericytes in the peri-infarct region across acute and chronic phases of ischemic stroke. Our results identify PSs as key regulators of microvascular perfusion during ischemia, with an excessive Ca²⁺ sensitivity and pathological contraction to ischemia and unique influence on adjacent contractile pericytes. We found that PS-adjacent contractile pericytes experienced severe Ca²⁺ increases and excessive vasoconstriction in the acute phase, leading to pericyte coverage loss, impaired vascular regulation, and compromised NVC responses in the downstream capillaries that persisted into the chronic phase. In contrast, non-PS-associated contractile pericytes exhibited more stable Ca²⁺ dynamics during the acute phase, therefore better preservation of coverage, and maintained vascular responses. Although pericyte coverage and Ca^2+^ signaling partially recovered from the second week post stroke, NVC responses remained suppressed. This segment-specific disparity in mural cell behavior appears to contribute to neurovascular uncoupling.

PSs have been identified as gatekeepers of capillary blood flow under physiological conditions^18^, while our study is the first to demonstrate their critical role during the progression of ischemic stroke. Earlier studies have shown that pericytes in the capillary bed contract during ischemia and hinder reperfusion, contributing to no-reflow after reperfusion. ^12,17^ Our data reveal that PSs are more contractile and more susceptible to ischemic Ca²⁺ rises than other mural cells in the ACT zone. The observed Ca^2+^ changes in pericytes adjacent to PSs may result from intercellular communication through direct membrane contacts such as gap junctions, as well as endothelial-mediated transmission^34–37^, allowing the propagation of vasoactive signals along the vascular wall in response to PS activity.

Consistent with previous studies^17,26,27^, we observed pericyte coverage loss during the chronic phase, which may result from excessive Ca^2+^ increase occurring in the acute phase. Ca^2+^ overload has been linked to calpain activation and mitochondrial or cytoskeletal injury in other injury contexts.^38–40^ Additionally, excessive intracellular Ca^2+^ can drive Ca²⁺ influx into mitochondria, elevating mitochondrial membrane potential, disrupting mitochondrial function, and triggering the caspase cascade, ultimately culminating in cell death. ^23–25^ Although we observed strong correlations between Ca²⁺ increases and cell death in both immunostaining and TPM imaging, the mechanisms remain obscure. Our acute-phase observations align with earlier reports^12,17,41,42^ that pericytes contract after ischemia. However, in contrast to studies describing persistent “die-in-rigor” phenotype, our data suggest that such constriction is largely confined to the acute phase. In the chronic peri-infarct region, we instead observed marked and prolonged capillary dilation in regions with unpreserved pericyte coverage (**Fig. 3J**), potentially reflecting loss of contractile regulation and, possibly, clearance of damaged pericytes by glial cells and macrophages.^43,44^ These findings suggest that pericyte contraction is a transient event, occurring only during the acute phase of ischemia. During the chronic phase, the peri-infarct region is characterized by localized capillary dilation and impaired NVC. These observations refine previous models by highlighting a temporal transition in microvascular behavior rather than a persistent constricted state.

Although SMCs, PSs, and contractile pericytes all exhibited excessive Ca^2+^ elevations during the acute phase, it was primarily contractile pericytes, particularly those on first-order capillaries, that demonstrated a higher reduction of pericyte coverage. Notably, PSs showed even greater Ca^2+^ increases than pericytes (PSs: 264.5 ± 42.31%; PAs: 260.9 ± 47.31%; first-order capillaries: 201.9 ± 30.79%; second-order capillaries: 142.6 ± 22.78%; third-order capillaries: 107.7 ± 37.36%) yet remained more structurally intact.

This apparent paradox suggests a dissociation between functional vulnerability and structural resilience. Functionally, PSs may be more vulnerable, exhibiting exaggerated Ca²⁺ elevations and strong constrictive responses that contribute to increased vascular resistance. In contrast, structurally, PSs appear more resistant to degeneration than downstream contractile pericytes. Several factors may contribute to this structural resilience. First, higher baseline Ca^2+^ levels in PSs and SMCs (**Fig. 1B**) may indicate an intrinsic tolerance to Ca^2+^ fluctuations during acute stress. Second, their robust cellular morphology, including thicker cell bodies and more circumferential vessel coverage^13,18,20,45^ (**Fig. 2B**; **Fig. 3C**), likely offers greater mechanical stability and resistance to ischemia compared to the thinner, sparser distributed pericytes. Third, the high expression of ɑ-SMA in PSs and PAs (**Fig. 2B**) may help maintain cytoskeletal integrity under pathological conditions.

Moreover, under healthy conditions, ACT regions containing PSs showed smaller vasodilatory responses and weaker mural cell Ca²⁺ responses to whisker air-puff stimulation (**Fig. 4; Supplemental Fig. 6**). Importantly, this should not be interpreted as reduced responsiveness of PSs themselves, which are known to exhibit strong contractile and vasoactive responses. Rather, our analysis captures pericytes within PS-associated ACT regions as integrated functional units. Within this context, pericytes downstream of PSs exhibited reduced Ca²⁺ responses and attenuated vasodilation, suggesting that the presence of a PS may constrain downstream vascular responsiveness.

This reduced baseline responsiveness may reflect a limited capacity to accommodate large Ca²⁺ fluctuations and may be associated with increased vulnerability during the acute phase, as ischemic stroke induced larger Ca²⁺ elevations in PS-containing ACT regions, where the majority of first-order capillaries subsequently exhibited pericyte coverage loss. As excessive intracellular Ca²⁺ may trigger apoptotic pathways^23–25,38^, the acute Ca²⁺ surge may contribute to the increased susceptibility to pericyte loss during the chronic phase. Future studies are needed to determine the Ca²⁺ threshold that triggers contractile pericyte loss.

The resulting contractile pericyte coverage loss leads to structural and functional remodeling of downstream capillaries, including chronic dilation and blunted reactivity, which collectively contribute to impaired perfusion adaptability during both baseline conditions and NVC responses (**Fig. 3**; **Fig. 5**). Neuronal loss post-stroke may partially contribute to the overall reduction in Ca^2+^ signaling and vascular responses observed in both regions with and without pericyte coverage loss. However, when capillaries were stratified based on pericyte presence, those lacking pericytes exhibited more pronounced impairment in vascular reactivity, along with a decoupling between Ca^2+^ dynamics and diameter changes. This spatially localized dysfunction is unlikely to be explained solely by impaired neuroglial signaling and instead suggests a direct consequence of pericyte coverage loss. As a result, these injury-prone regions exhibit a failure to adequately recruit blood during functional hyperemia, further exacerbating neurovascular uncoupling and tissue hypoperfusion.

The heterogeneity of pericyte dysfunction following ischemia disrupts the spatial and temporal coherence of capillary blood flow regulation (**Fig. 6**). In areas where pericytes remain functional, blood can still flow to hungry neurons after 7 days post-stroke (**Fig. 6F, G**). In contrast, regions with pericyte dysfunction experience delayed and weak responses, resulting in spatial displacement of NVC responses. This mismatch between perfusion and metabolic demand leads to further impaired functional hyperemia. Notably, although structural recovery of contractile pericytes partially restored basal vascular tone, it was insufficient to restore normal neurovascular coupling after ischemic injury (**Fig. 7**). Those persistent NVC impairments can worsen the neurological outcome and slow functional recovery following stroke^3^. This differential vulnerability underscores the importance of considering microvascular heterogeneity in stroke pathophysiology.

Several limitations of this study should be acknowledged. First, while TPM provides high-resolution data on Ca²⁺ dynamics, our measurements are limited to cortical surface microvessels and may not fully capture deeper or regional heterogeneity. Second, although we observed strong correlations between acute Ca²⁺ surge, contractile responses, and pericyte coverage loss, causal mechanisms require further validation using genetic or pharmacological manipulation of Ca²⁺ handling. Finally, our study was conducted in a rodent model of stroke, and future research is needed to confirm the relevance of these mechanisms in human tissue. Future studies should aim to dissect the upstream triggers of Ca²⁺ dysregulation in PSs, explore their interactions with endothelial and immune cells during stroke progression, and test therapeutic interventions that preserve PS function. It will also be important to determine whether PS dysfunction is reversible or represents a point of no return in stroke-induced microvascular damage.

In conclusion, our findings identify PSs as a functionally distinct and highly vulnerable microvascular component during ischemic stroke. Through their abnormal Ca²⁺-dependent contractility and profound influence on surrounding mural cells, PSs contribute to impaired reperfusion, disrupted neurovascular coupling, and chronic capillary dysfunction. Targeting these mechanisms may offer a better recovery after traditional large-vessel recanalization approaches.

## Methods

### Animal

Acta2-GCaMP8.1/mVermilion (Tg (Acta2-GCaMP8.1/mVermilion) B34-4Mik, JAX #032887; Jackson Laboratory) mice and C57BL/6J mice (Jackson Laboratory) of both sexes, aged 10 to 22 weeks, were used in this study (12 males and 10 females). All procedures were approved by the Danish National Committee on Health Research and conducted in accordance with the European Council’s Convention for the Protection of Vertebrate Animals used for experimental and other scientific purposes, adhering to established animal research guidelines. All mice were maintained under a 12-hour light/dark cycle, with ad libitum access to food and water via the cage lid.

A total of 22 mice were used in this study. Cohorts were allocated as follows: acute two-photon imaging (N = 5), chronic awake two-photon imaging (N = 8), sham longitudinal imaging (N = 4), and immunohistochemistry (N = 5). LSCI analyses were performed in a subset of the chronic imaging cohort (N = 7). Cohorts were partially overlapping, with some mice contributing to multiple experimental paradigms.

### Chronic Cranial Window Implantation

Mice were anesthetized with 1–2% isoflurane and placed in a stereotaxic frame on a temperature-controlled heating pad (37 °C), with body temperature continuously monitored via a rectal probe. Preoperative analgesia and anti-inflammatory treatment included subcutaneous carprofen (5 mg/kg), buprenorphine (0.05 mg/kg), dexamethasone (4.8 mg/kg), and 0.5 mL sterile saline; 0.1 mL Lidocaine was injected locally at the surgical site.

A cranial window (Ø ∼4 mm) was drilled over the left hemisphere (anterior-posterior: –1.5 mm; medial-lateral: −3 mm), covered with a 5 mm glass coverslip, and secured using 3M™ RelyX™ resin cement. A lightweight stainless steel head plate (Neurotar®) was affixed to the skull using Paladur® dental cement, ensuring complete coverage with no exposed skull.

Postoperatively, mice received 0.5 mL sterile saline and were maintained on a heating pad overnight. Analgesia (carprofen and buprenorphine) continued for 3 days. Animals were monitored daily for 7 days, and experiments were performed ≥ 4 weeks after surgery (**Fig. 1A**).

### Transient Filament Middle Cerebral Artery Occlusion

Transient MCAO was performed as described by Longa et al.^46^. Mice were anesthetized with 1−2% isoflurane and a silicone-coated monofilament was introduced via the external carotid artery into the internal carotid artery to occlude the middle cerebral artery. Occlusion was maintained for 30 min (chronic experiments) or 60 min (acute experiments), followed by reperfusion through filament withdrawal. Sham-operated animals underwent identical surgical procedures without occlusion of middle cerebral artery. Successful occlusion was verified by laser Doppler flowmetry.

Preoperative analgesia included subcutaneous carprofen (5 mg/kg), buprenorphine (0.05 mg/kg), and 0.5 mL sterile saline. Filament size was selected based on body weight to ensure consistent occlusion. A 7-0 silicone-coated monofilament (coating diameter 0.17 ± 0.01 mm; coating length 4–5 mm; Doccol Corporation, USA) was used for mice weighing <30 g, whereas a 0.19 mm tip diameter filament was used for weighing ≥30 g.

Neurological deficits were assessed using the extended neurological scoring system (**Supplemental Fig. 1B**), as described by Garcia *et al*.^47^. Postoperative supportive care, including daily subcutaneous administration of carprofen, buprenorphine, and saline, was provided for three days to minimize mortality. Body weight and general behavior were monitored daily during the first postoperative week.

### *In Vivo* Imaging During the Acute Phase of Ischemic Stroke

Acta2-GCaMP8.1/mVermilion mice were anaesthetized with 1% isoflurane, with body temperature maintained as described above. The vascular lumen was labelled by retro-orbital injection of 50 μL 2% (wt/vol) Texas Red–dextran (70 kDa, Invitrogen).

Imaging was performed using a commercial TPM (FluoView FVMPE-RS, Olympus). Epifluorescence images were acquired using a 4×, 0.16 NA air objective and a scientific CMOS camera (ORCA-Flash 2.8, Hamamatsu). Capillary segments, including PS-associated and non−PS-associated regions, were mapped using a 25×, 1.05 NA water-immersion objective and identified on epifluorescence angiograms (**Supplemental Fig. 1**).

Mural cell Ca^2+^ dynamics and vascular diameters were collected before MCAO, 5 min after insertion of filament, and 10–15 min after the reperfusion. Data were acquired based on epifluorescence angiograms using volumetric 4D (x–y–z–t) hyperstack imaging with the 25×, 1.05 NA objective, covering vascular segments from PAs to second- or third-order capillaries. Image stacks were acquired at a spatial resolution of 0.118–0.495 µm/pixel (512 × 512 pixels), with a z-step of 2.0–2.58 µm, comprising 11–27 planes per stack, at a scan speed of 2.0 µs/pixel (∼15 s per stack). Each ROI was recorded for six imaging cycles and averaged for analysis.

For ROIs containing PSs, high-resolution imaging was performed. Stacks were acquired at 0.026–0.062 µm/pixel (1024 × 1024 pixels), with a z-step of 1.5 µm, comprising 6–17 planes per stack, at 2.0 µs/pixel (15–32 s per stack). Each ROI was recorded for three cycles and averaged.

Excitation wavelength was set to 920 nm. Emission from GCaMP8.1 and Texas Red–dextran was collected using 489–531 nm and 601–657 nm filters, respectively, with identical laser settings across time points.

### *In Vivo* NVC Imaging During Chronic Phase of Ischemic Stroke

Longitudinal, awake imaging was performed using a commercial TPM (Femto3D-RC, Femtonics) equipped with a 25×, 1.0 NA piezo-driven objective and controlled by Femtonics MES v6 software. Recordings were conducted at days 0 (baseline), 1, 4, 7, 14, and ≥21 post-stroke (**Fig. 3A, B**).

Fast repetitive hyperstack imaging (4D imaging) was performed for each ROI (**Fig. 3B**), covering segments from PAs to second-order capillaries. This approach enabled the visualization of entire z-axis ranges of the targeted vessels at desired spatial and temporal resolution. Each image stack was acquired in under 1 second and consisted of 6–11 planes with a step size of 3.5–5.5 µm. Pixel size in the x–y plane ranged from 0.33 to 0.57 µm/ pixel. Excitation was provided at 920 nm, and emitted fluorescence was spectrally separated to capture green signals from GCaMP8.1-expressing Acta2-positive mural cells and red signals from Texas Red-dextran-labeled vessel lumens.

During awake NVC imaging, whisker air-puff stimulation was delivered during hyperstack acquisition, following a baseline period of at least 30 s and 60 s post stimulation. Air-puff stimulation (1 s duration at 3 Hz, 0.25 s pulse width) was applied to the distal whiskers via a metal tube. Facial behaviors (pupillometry and whisking) and body behaviors (locomotion and grooming) were monitored using two synchronized cameras. Real-time locomotion speed was recorded using the integrated Neurotar Mobile HomeCage magnetic tracking system, which computes cage position and velocity from magnetic sensors independent of lighting conditions (**Fig. 3B**). Pupil size, whisking, and locomotion were recorded simultaneously with TPM.

Trials exhibiting movement artifacts (locomotion speed >1 cm/s or overt limb activity such as grooming or lifting) were excluded from analysis. Only motion-free trials were included in whisker-evoked response analyses. For each mouse, data were collected from three ROIs.

### Laser Speckle Contrast Imaging

Cortical cerebral blood flow was measured using LSCI in awake Acta2-GCaMP8 mice immediately following TPM at days 0 (baseline), 1, 4, 7, and 14 post-stroke (**Fig. 3A**). The recording setup was customized as previously described.^48^ Illumination was provided by a volume holographic grating–stabilized laser diode (LP785-SAV50, Thorlabs) controlled by a CLD1011LP driver (Thorlabs). Images were acquired using a CMOS camera (acA2040-90umNIR, Basler) coupled to a 5× objective (0.15 NA; HCX PL FLUOTAR, Leica), yielding a spatial resolution of 3.19 µm/pixel (1024 × 1024 pixels) at 50 frames per second.

Whisker stimulation protocols were identical to those described above. To minimize stress, total recording time per mouse per session was limited to 1.5 h (1.25 h TPM and 15 min LSCI). Sessions were terminated early if signs of distress were observed.

All LSCI data were processed offline using custom MATLAB (R2021b) scripts developed based on previously published method.^49^ Briefly, spatial contrast *K* was calculated for each pixel as the ratio of the standard deviation to the mean of the intensities within the 5×5-pixel neighborhood, and the blood flow index (BFI) was subsequently computed as BFI = 1/K^2^ to obtain perfusion dynamics.

### Two-Photon Imaging Analyses

All data were processed using ImageJ/Fiji or custom-written scripts developed in MATLAB (R2021b).

#### Identification of Precapillary Sphincters

As described in a previous study^18^, the PS is defined as a mural cell encircling an indentation of the capillary at the point where it branches from a PA. In this study, PSs were identified based on the following three criteria: (1) Location: situated at the junction between the PA and a first-order capillary; (2) Diameter: the indentation at the junction was < 0.8× the diameter of the first-order capillary; and (3) Morphology: the indentation was encircled by a thick mural cell (GCaMP8.1 expression). Only regions fulfilling all three criteria were classified as PSs.

#### Pericyte Coverage Change Identification and Classification

For longitudinal analyses, each pericyte ROI was tracked across time and classified according to its signal state and fate. All preprocessing steps were kept identical across time points within each mouse.

Pericyte Ca^2+^ signals were quantified within predefined vascular ROIs using ImageJ/Fiji on calibrated images. Signal segmentation within each ROI was performed using ImageJ’s *Auto Threshold* function. The thresholding algorithm was selected at baseline (D0) for each ROI based on segmentation quality and separation of pericyte signal from background, and the same algorithm was applied consistently to that ROI at all subsequent time points. Thresholds were restricted to the ROI to avoid bias from unrelated bright structures outside the mural cells.

#### Quantification of coverage and signal intensity

Pericyte coverage was defined as the fraction of ROI pixels above the selected threshold. To determine whether apparent coverage loss reflected true signal disappearance rather than poor signal-to-noise ratio or residual weak fluorescence, intensity metrics were further quantified. Within each ROI, the mean intensity of pixels above (mean_high_) and below (mean_low_) threshold were quantified. Background mean (μ_bg_) and standard deviation (σ_bg_) were measured from a fixed parenchymal background ROI for each mouse and used to compute Z-scores (**Supplemental Fig. 4A**):

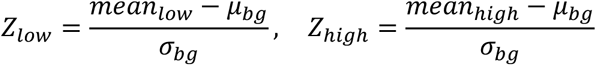

#### Classification of pericyte’s preservation state

ROIs were classified into two states: unpreserved or preserved pericytes. An ROI was classified as unpreserved if all the following criteria were met (**Supplemental Fig. 4A-H**):

1. Reduced Ca^2+^ coverage: threshold coverage decreased to < 50% of baseline (D0).
2. Loss of local signal: Signal below the threshold was indistinguishable from background (mean_low_ < μ_bg_ + 3× σ_bg_; Z_low_ < 3).
3. Adequate signal-to-noise: Signal above the threshold in remaining regions exceeded background (mean_high_> μ_bg_ + 4×σ_bg_; Z_high_ > 4), ensuring signal loss was not attributable to global image degradation.

To reduce classification driven by transient fluctuations, these criteria were required to be satisfied for at least two consecutive imaging sessions or at the terminal time point. ROIs not fulfilling all criteria were classified as preserved.

#### Identification of pericyte regrowth

ROIs classified as unpreserved were subsequently assessed for restoration. An unpreserved ROI was classified as a restored state if all the following criteria were met (**Supplemental Fig. 4D, H**):

1. Restored coverage: threshold coverage increased to ≥ 50% of baseline or > min prior coverage + 20%
2. Exclusion of background-driven coverage increases: Mean intensity of signal below the threshold was higher than at least the level observed at the time of min prior coverage (Z_low_ ≥ Z_low___min prior coverage_), ensuring that increased coverage was not due to background fluorescence being misclassified as signal.
3. Adequate signal-to-noise: Mean suprathreshold intensity remained well above background (mean_high_> μ_bg_ + 4×σ_bg_; Z_high_ > 4), ensuring that apparent restoration was not driven by imaging noise or global signal fluctuations.

This combined intensity-based, coverage-based, and structural approach enabled robust discrimination between true pericyte signal un-preservation and apparent reductions arising from weak fluorescence, reduced signal-to-noise ratio, or threshold-dependent segmentation artifacts.

#### Analysis of data in the acute phase of stroke

During the acute phase, vessel diameters and mural cell Ca^2+^ intensities were quantified from TPM generated by average-intensity projection using ImageJ/Fiji. Vessel diameters were measured by placing multiple linear ROIs perpendicular to the vessel lumen and averaging multiple measurements across ROIs at each capillary order. For Ca^2+^ signal analysis, ROIs were drawn along the morphological boundaries of mural cells at different vascular segments. Default thresholding method was applied to delineate GCaMP8.1-labeled mural cells in the image by mean intensity projection across the whole video, and only pixels above threshold were included in the analysis. ROIs were placed at identical locations and in the same number across time points, using mural cell morphology as a reference.

#### Whisker air puff stimulation analysis in the chronic phase of stroke

For fast repetitive hyperstack imaging during whisker stimulation, image stacks were first motion-corrected and then collapsed along the z-axis using average-intensity projection to generate a time series of projected images (x–y–t), which was used for all subsequent analyses (**Fig. 3B**). Vascular diameter responses were quantified from the projected image series using a custom MATLAB algorithm as described before.^20,48^ Rectangular ROIs (2–4 µm wide) were placed perpendicular to the vessel axis at predefined locations. Vessel boundaries were detected over time using a Chan–Vese active contour segmentation algorithm, and diameter changes were extracted accordingly.

Ca^2+^ responses to whisker stimulation were quantified from the same projected image series using ImageJ/Fiji, following the protocol used for acute-phase analysis. ROIs were placed at identical locations and in the same number across time points, using mural cell morphology as a reference.

#### Resistance calculation in the arteriole-capillary transitional (ACT) zone

To estimate the change in flow resistance upon changes in vessel diameter of all segments in the ACT zone. We adapted Poiseuille’s law (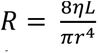, where R is resistance, η is blood viscosity, L is vessel length, r is vessel radius), neglected constants (η) and fixed numbers. We therefore simplified the equation as 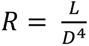 (where L is vessel length and D is vessel diameter), to individually calculate resistance at PAs, PSs, first- to third-order capillaries at baseline, occlusion and reperfusion stages. Next, we used the series-parallel circuit configuration shown in **Fig. 1Q** to calculate the total resistance in the ACT zone with or without PSs. The equation is given below, with *R*_*s*p*h*_ included only when a PS was present

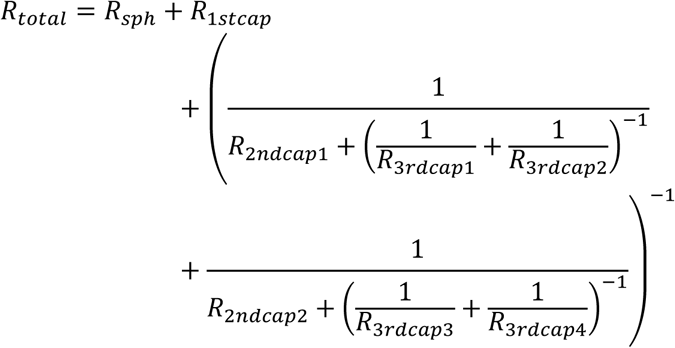

In a limited number of cases, diameter measurements were not available for all four third-order capillaries. In these cases, missing values were approximated using diameter measurements from third-order capillaries within the same branching pathway from the second-order capillary.

### Brain Collection

C57BL/6J mice underwent MCAO surgery and postoperative care as described before, with an occlusion duration of 30 minutes. Four days after MCAO, mice deeply anesthetized with xylazine (10 mg/kg) and ketamine (100 mg/kg) administered intraperitoneally, and transcardially perfused at a rate of 10 mL/min with 0.1 M phosphate-buffered saline (PBS) for 2−3 minutes, followed by 4% paraformaldehyde in 0.1 M PBS for 4−5 minutes. Brains were extracted and post-fixed in 4% paraformaldehyde at 4 °C for 2 hours, then transferred to 30% sucrose at 4 °C for 48 hours. Samples were subsequently frozen in dry ice-chilled isopentane and stored at approximately –80 °C until use.

### Immunohistochemistry and Confocal Imaging

Brains were sectioned coronally at 50 μm thickness using a cryostat (Leica CM3050 S). Sections were rinsed in PBS for 5 minutes, three times. Permeabilization and blocking were performed overnight at 4 °C in PBS containing 1% bovine serum albumin and 0.5% Triton X-100. Sections were then incubated at 4 °C for two nights with the following primary antibodies: rat anti-podocalyxin (1:400, MAB1556, R&D Systems), goat anti-PDGFR-β (1:250, AF1042, R&D Systems), and mouse anti-α-SMA-FITC (1:200, F3777, MilliporeSigma). After three 5-minute PBS washes, sections were incubated overnight at 4°C with secondary antibodies: donkey anti-goat IgG H&L (Alexa Fluor 568) (1:500, A-11057, Invitrogen), donkey anti-rat IgG H&L (Alexa Fluor 647) (1:500, A78947, Invitrogen). Following three additional 5-minute PBS washes, sections were stained with Hoechst 33342 (1:6000, B2261, Sigma) for 7 minutes. After a final series of three PBS washes, sections were mounted using SlowFade Diamond antifade mountant (S36963, Invitrogen) and sealed with nail polish.

Stained sections were scanned using Zeiss Axioscan 7 with a 20× objective at a resolution of approximately 0.172 μm/pixel (x and y) and 3 μm (z-step) for whole-slice overview imaging. High-resolution images were acquired using a Zeiss LSM 980 confocal microscope equipped with a 40× water-immersion objective.

Image analysis was performed by FiJi (US National Institutes of Health). The distance from PAs to the endpoint of contractile pericyte coverage, as well as the capillary branch order at the endpoint, were quantified based on α-SMA (marking VSMCs and contractile pericytes) and podocalyxin (marking vessels) expression.

### Statistical Analyses

Correlation analyses were conducted using custom MATLAB scripts (R2021b). Frequentist statistical analyses were performed using R Studio, and Bayesian statistical analyses were implemented in Python using custom-written code.

#### Correlation analyses between diameter and calcium

To assess the coupling between vascular diameter changes and Ca^2+^ dynamics at different stages following ischemic stroke, correlation analyses were performed separately for acute and chronic phases. During the acute phase, Ca²⁺ changes were tested for correlation with corresponding changes in vessel diameter. During the chronic phase (whisker stimulation), Ca²⁺ responses were tested for correlation with the temporal derivative of vessel diameter to capture dynamic vascular responses. Pearson correlation coefficients and regression slopes were calculated to quantify the relationship between these measures. Correlations and regression slopes were computed separately for each ROI and subsequently averaged per mouse for group-level statistical comparisons. Statistical significance was defined as *P* < 0.05.

#### Frequentist statistical analyses

Data are presented as mean ±SEM, with individual data points shown where possible. Unless otherwise stated, the inferential unit was the mouse (N = number of mice). ROI counts are reported to describe sampling density, but statistical testing was performed on mouse-level summaries or using linear mixed-effects models that accounted for clustering of repeated observations within mice. Paired *t*-tests were used for comparisons between two paired groups. Linear mixed-effects models were fit in R and R studio to account for clustering/repeated measures (e.g., random intercepts for mouse and, where relevant, vascular location nested within mouse). Fixed effects (e.g., PS-associated or non−PS-associated, preserved or unpreserved pericytes, time points post stroke, and their interaction) were tested using Type III inference with Satterthwaite’s approximation for denominator degrees of freedom, reporting two-sided p values. Statistical significance was defined as * *P* < 0.05, ** P < 0.01, *** *P* < 0.001, and **** *P* < 0.0001.

#### Bayesian state space analysis of time-dependent vascular changes

For data measured across more than three time points (**Supplemental Fig. 7B–I, K–N; Supplemental Fig. 9B–I**), a Bayesian state space modelling approach was used to account for temporal dependence between repeated measurements. This framework allows estimation of time-varying effects while accommodating measurement noise and irregular sampling intervals, following a previously published approach for microvascular dynamics^19^.

Temporal dynamics were modeled using a Gaussian random walk prior. The latent standardized response 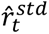 evolved as:

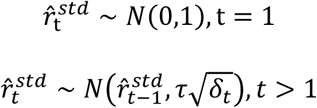

where *δ*denotes the time interval between observations and *τ* controls temporal variability.

Weakly informative priors were used throughout. Posterior inference was performed using adaptive Hamiltonian Monte Carlo sampling with the Python library PyMC^50^. Model convergence was assessed using standard diagnostics, including improved R̂ convergence statistic^51^, effective sample sizes, and the absence of post-warmup divergent transitions. Model adequacy was evaluated via posterior predictive checks.

#### Bayesian analysis of capillary order measurements from immunochemistry

Capillary order at the endpoint of α-SMA transition (values 1–6) was analyzed using a Bayesian ordinal regression model (**Fig. 2D, G**), preserving its ordered structure without assuming equal spacing. The model included capillary type (PS-associated vs. non−PS-associated), anatomical site (ipsilateral vs. contralateral), their interaction, and a random intercept for mouse:

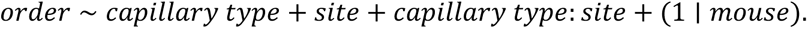

The model was implemented and fitted using Markov chain Monte Carlo sampling with the Python library Bambi^52^, using weakly informative prior distributions. Convergence was assessed using standard diagnostics.

Statistical conclusions were based on posterior distributions rather than null hypothesis significance testing. For each comparison, we calculated the posterior probability that one condition exceeded the other; probabilities > 0.95 (or < 0.05) were considered strong evidence for a directional effect. Code is available at: https://github.com/teddygroves/stroke/.

## Data Availability

The data that support the findings of this study and custom analysis code are available from the corresponding author upon reasonable request.

## Code availability

The custom analysis code used in this study is available at: https://github.com/teddygroves/stroke/.

Correspondence and requests for materials should be addressed to Changsi Cai.

## Acknowledgement

We thank Micael Lønstrup for training and assistance with microsurgery. We thank the technical assistance of the Core Facility for Integrated Microscopy, University of Copenhagen. We used ChatGPT for grammar checking and language refinement during manuscript preparation. Some images were created using Biorender.com under license. This study was supported by Lundbeck Foundation (#R345-2020-1440, #R436-2023-1125, #R392-2018-2266, and #R345-2020-1440), the Independent Research Fund Denmark (#1133-00016B and #1030-00374A), the Novo Nordisk Foundation (#0064289, #0092323, #117272, #14CC0001), EU Horizon Europe (#101060066), Læge Sofus Carl Emil Friis og Hustru Olga Doris Friis’ Legat, Dagmar Marshalls Fond, Familien Hede Nielsens Fond, Hørslev fonden, Oda og Hans Svenningsens Fond, Helsefonden, and ZonMw VENI #09150162410015 (to IAM).

