## Supplemental Fig. 1-10 for "Impaired Calcium Signaling in Precapillary Sphincters and Pericytes Perturbs Neurovascular Regulation after an Ischemic Stroke"

### Supplementary materials

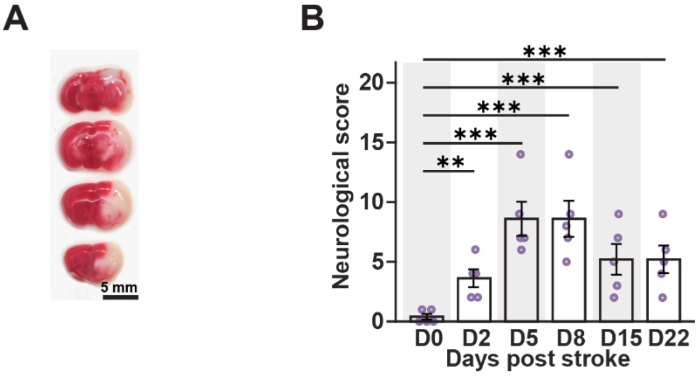


**Supplemental Figure 1. Neurological scores of Acta2-GCaMP8 mice across different days and infarct core.** (A) Representative 2,3,5-triphenyltetrazolium chloride staining from a mouse subjected to middle cerebral artery occlusion (MCAO) (scale bar: 5 mm).
(B) Comparison of neurological scores between baseline (D0) and different days post-stroke (N = 5 mice). Data are presented as mean ± SEM. Statistical analysis was performed using a linear mixed-effects model, with neurological scores treated as approximately continuous variables. * $P < 0.05$, ** $P < 0.01$, *** $P < 0.001$.

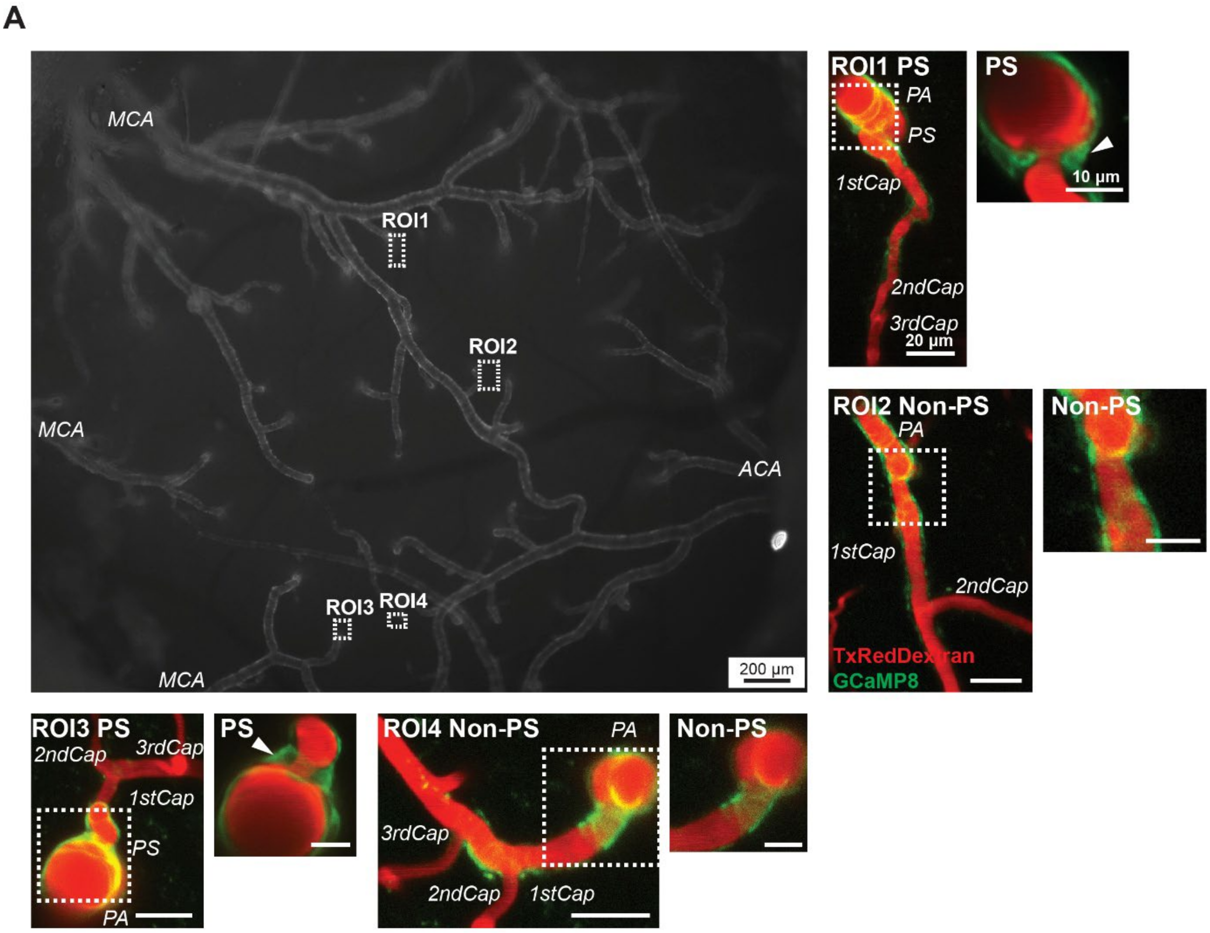


**Supplemental Figure 2 Mapping of PS-associated and non−PS-associated regions across the cranial window.** (A) Representative cortical vasculature across the entire cranial window (scale bar: 200 µm). In each mouse, one to two ROIs containing precapillary sphincters (PSs; PS-associated) and one to two ROIs lacking PSs (non−PS-associated) were selected and localized using pial arteries as anatomical references (white dashed squares). Zoomed-in views show that ROIs in the non−PS-associated group comprised penetrating arterioles (PAs) and first- (1stCap) to third-order capillaries (3rdCap), whereas ROIs in the PS-associated group comprised PAs, PSs, and first- to third-order capillaries (scale bar: 20 µm). Higher-magnification images highlight the morphology and anatomical location of PSs (scale bar: 10 µm). Arrowheads indicate PSs. Vessels are labeled with Texas Red–dextran (red), and mural cell $Ca^{2+}$ signals are visualized by GCaMP8 fluorescence (green). MCA: middle cerebral artery. ACA: anterior cerebral artery.

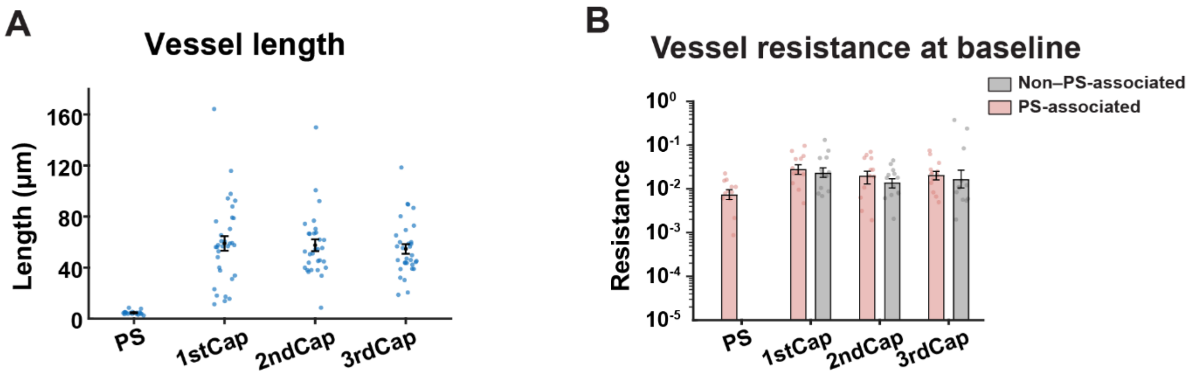


| Capillary Order | Mean Vessel Length (μm) |
|---|---|
| Sphincter | 4.64 ± 0.33 |
| 1stCap | 58.93 ± 5.64 |
| 2ndCap | 57.52 ± 4.64 |
| 3rdCap | 54.63 ± 3.79 |

**Supplemental Figure 3 Vessel length and resistance at baseline.**

(A) Vessel length across different vessel segments (Top: dot plot, n = 19−33 ROIs from 6 mice; bottom: mean ± SEM).

(B) Vessel resistance at baseline across different vessel segments in the non−PS-associated and PS-associated groups (n = 12 ROIs from 5 mice).

Data are mean ± SEM.

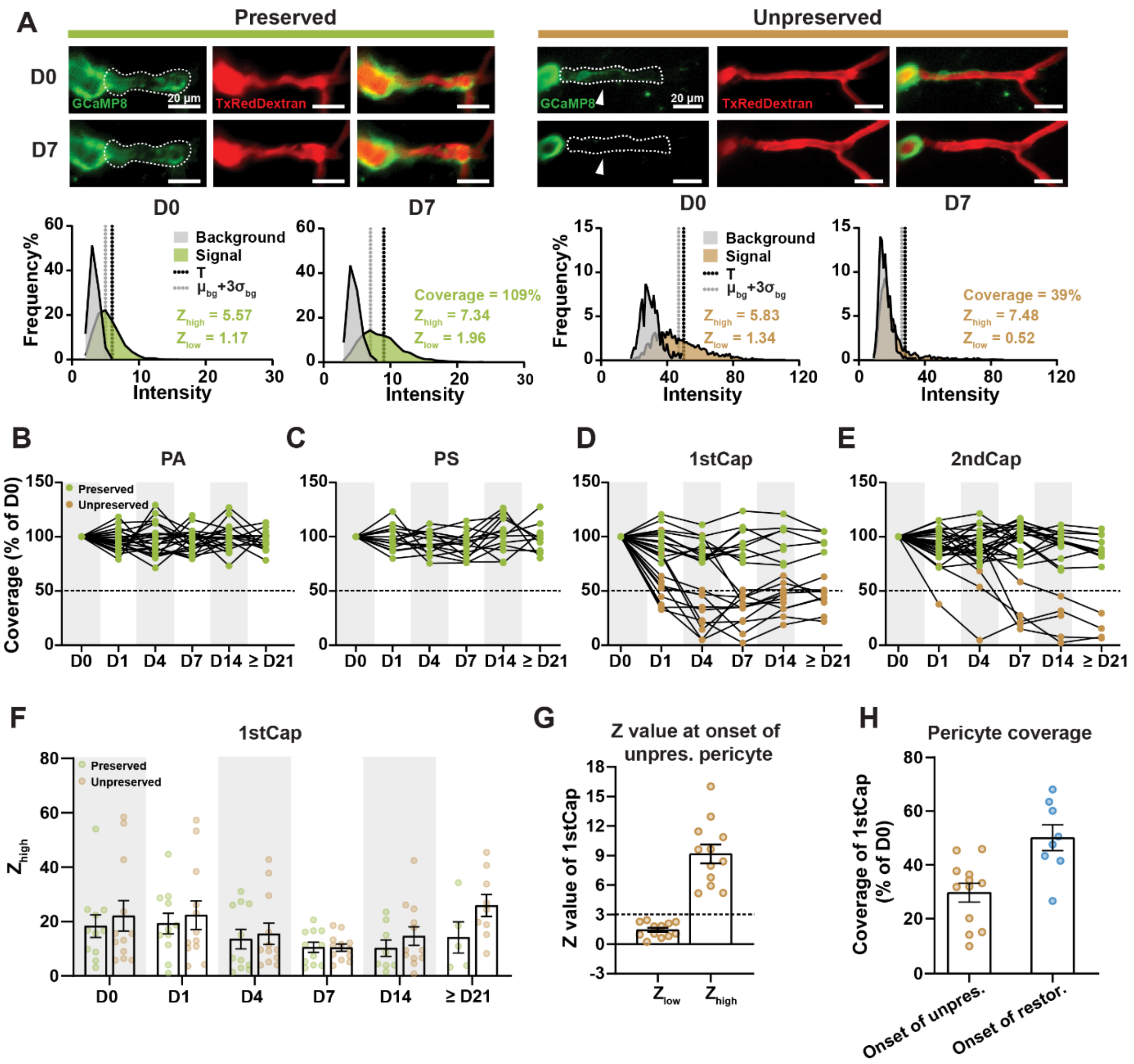


**Supplemental Figure 4. Identification and classification of pericyte coverage loss.**

(A) Representative images of preserved and unpreserved pericytes (top) and corresponding fluorescence intensity distributions in first-order capillaries and background regions (bottom) at baseline (D0) and day 7 post-stroke (D7) (scale bar: 20 µm). Vessels are labeled with Texas Red–dextran (red), and mural cell $Ca^{2+}$ signals are visualized by GCaMP8 fluorescence (green). For mural cells surrounding each vessel segment, $Ca^{2+}$ coverage relative to D0, as well as $Z_{high}$ and $Z_{low}$, were calculated. An ROI was classified as unpreserved if all of the following criteria were met: (1) Reduced $Ca^{2+}$ coverage, defined as high-threshold coverage < 0.5 of D0; (2) Loss of local signal, defined as mean intensity at the coverage-reduced site $< \mu_{bg} + 3\times \sigma_{bg}$ ($Z_{low} < 3$); and (3) Adequate signal-to-noise, defined as mean intensity of maintained regions $> \mu_{bg} + 4\times\sigma_{bg}$ ( $Z_{high} > 4$), ensuring sufficient imaging quality. Classification required confirmation in at least two consecutive imaging sessions or at the terminal time point. ROIs not fulfilling all criteria were classified as preserved.

(B−E) The percentage of coverage in PAs (B), PSs (C), first-order (D) and second-order (E) capillaries across different post-stroke days. Green indicates preserved ROIs, and brown indicates unpreserved ROIs. Data represent n = 14−23 ROIs from 8 mice in PAs, n = 10−15 ROIs from 8 mice in PSs, n = 5−12 ROIs from 7 mice in first-order capillaries, and n = 4−23 ROIs from 8 mice in second-order capillaries.

(F) $Z_{high}$ value in first-order capillaries of preserved and unpreserved groups across different post-stroke days. Green indicates preserved ROIs, and brown indicates unpreserved ROIs (n = 5−12 ROIs from 7 mice).

(G) $Z_{low}$ and $Z_{high}$ values of first-order capillaries at the onset of coverage loss (n = 12 ROIs from 7 mice).

(H) Pericyte coverage in first-order capillaries at the onset of coverage loss and restoration (n = 8−12 ROIs from 7 mice).

Data in F−H are presented as mean ± SEM.

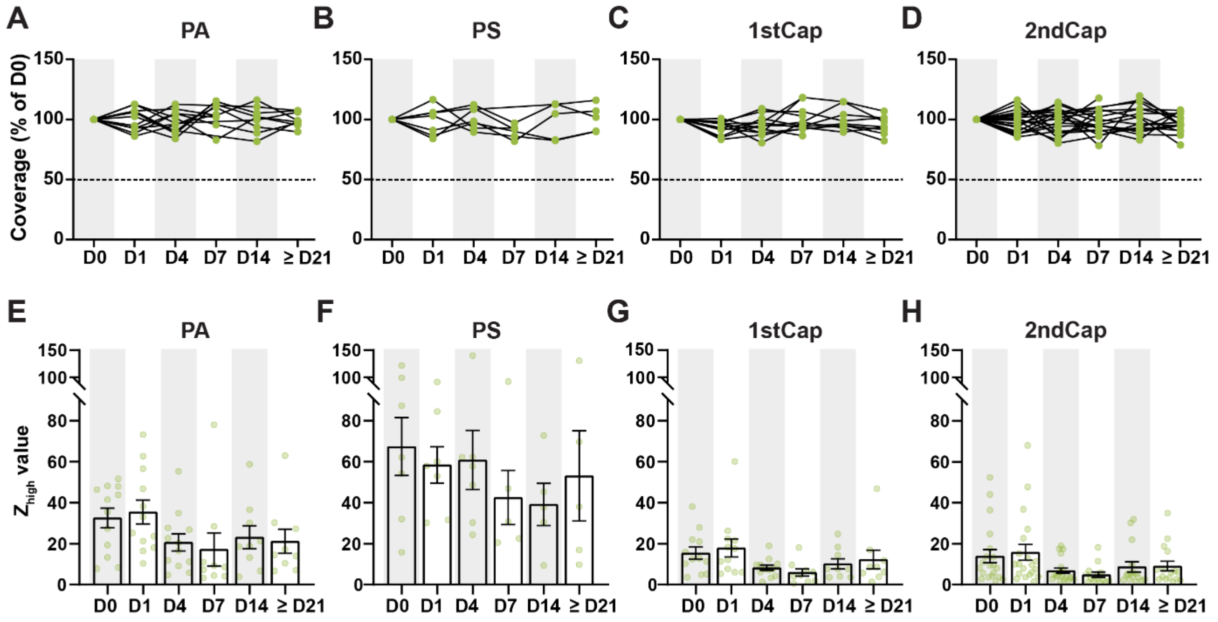


**Supplemental Figure 5. Pericyte coverage and $Z_{high}$ in sham mice across imaging sessions.**
(A−D) The percentage of coverage in PAs (A), PSs (B), first- (C) and second-order (D) capillaries across different post-stroke days.
(E−H) $Z_{high}$ value in PAs (E), PSs (F), first- (G) and second-order (H) capillaries across different post-stroke days. Data in E−H are presented as mean ± SEM. Data represent n = 9−12 ROIs from 4 mice in PAs, and first-order capillaries, n = 5−7 ROIs from 4 mice in PSs, and n = 14−20 ROIs from 4 mice in second-order capillaries.

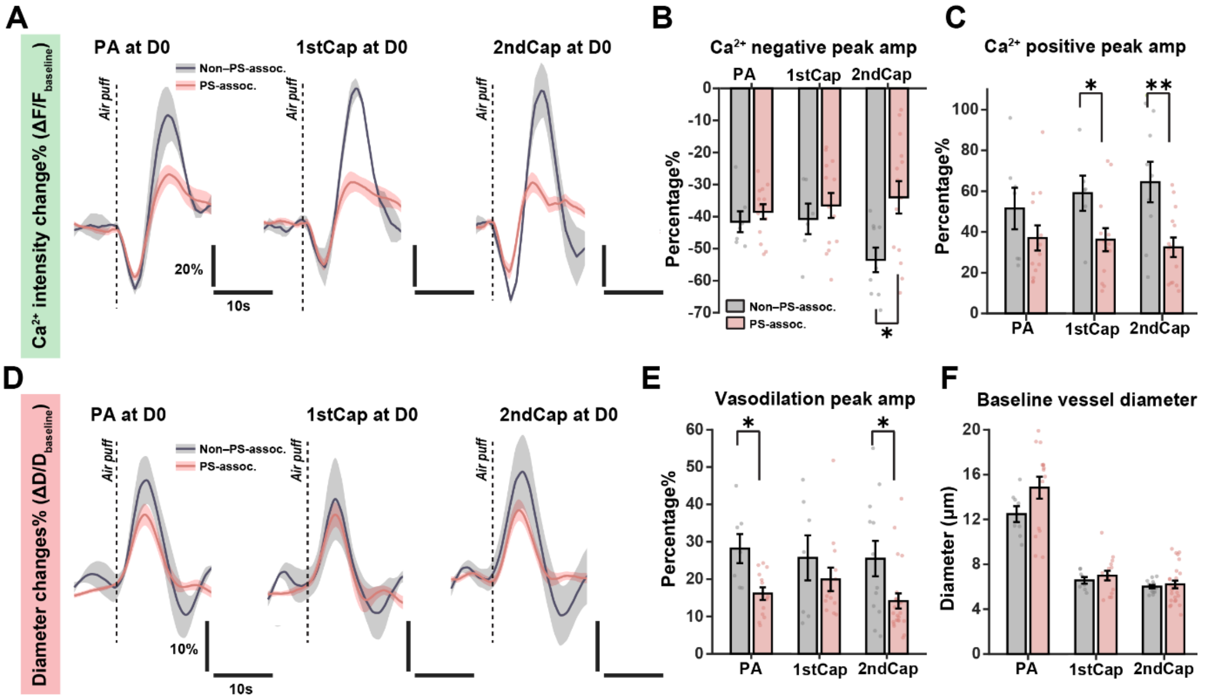


**Supplemental Figure 6. Calcium and vascular responses in PS-associated and non−PS-associated groups at D0.**

(A, D) $Ca^{2+}$ (A) and vascular (D) response curves in different mural cells and corresponding vessel segments in the PS-associated (pink) and non−PS-associated (grey) groups at D0. Each curve represents the mean response averaged across eight mice.

(B, C) Amplitude of negative (B) and positive (C) $Ca^{2+}$ peaks in PAs, first- and second-order capillaries at D0 in the PS-associated (pink) and non−PS-associated (grey) groups, normalized to the pre-stimulation baseline (25 s before air-puff induced whisker stimulation; N = 8 mice).

(E) Amplitude of peak vascular responses to whisker stimulation in PAs, first- and second-order capillaries at D0, normalized to the pre-stimulation baseline (25 s before air-puff induced whisker stimulation; N = 8 mice).

(F) Baseline vessel diameter in PAs, first- and second-order capillaries at D0 (N = 8 mice).

Data are presented as mean ± SEM. A linear mixed-effects model was applied for panels B, C, E, and F. Comparisons between PS-associated and non−PS-associated groups are indicated by * $P < 0.05$, and ** $P < 0.01$.

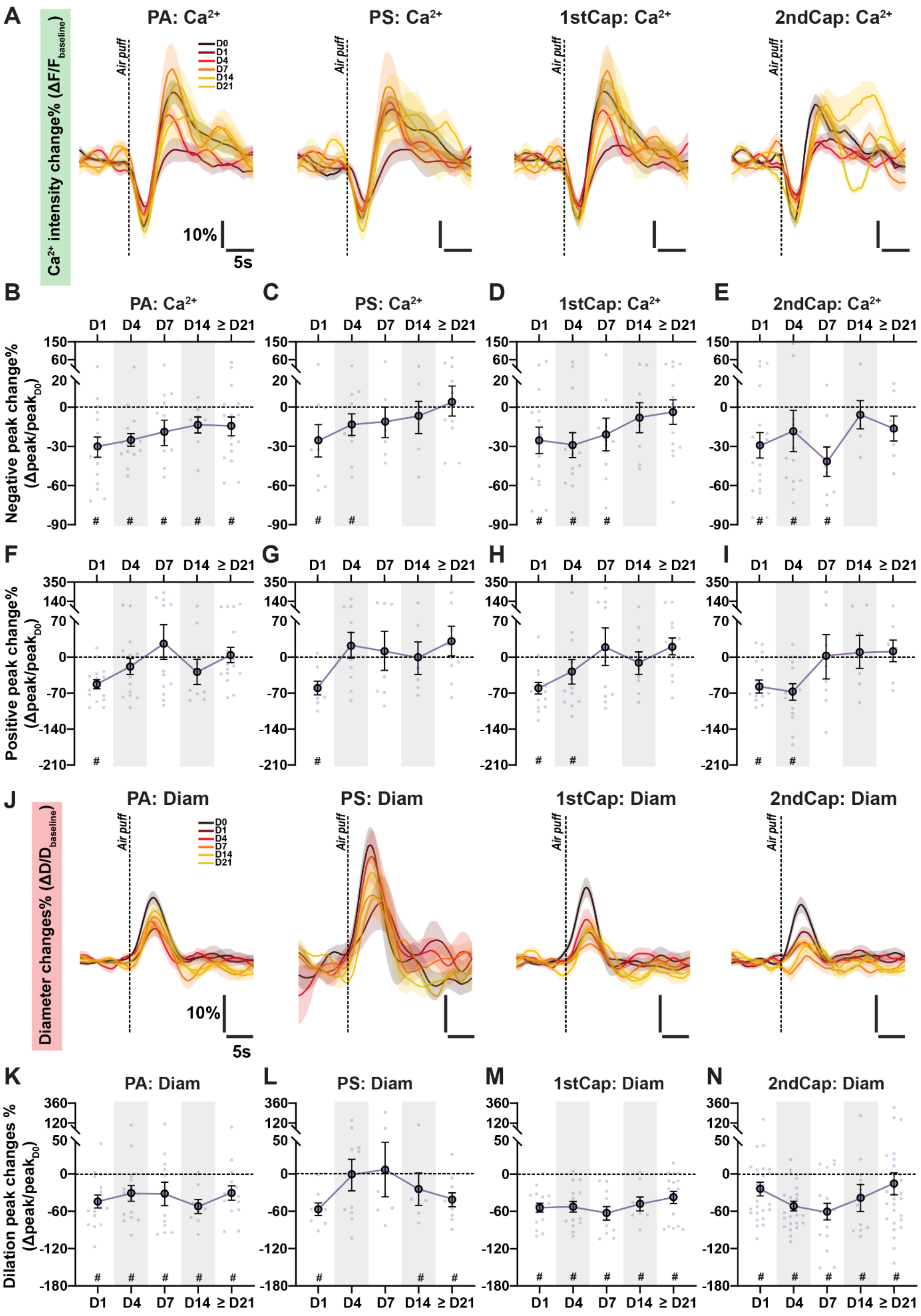


**Supplemental Figure 7. Ischemia reduces mural cell calcium and vascular responses to whisker stimulation.**

(A, J) $Ca^{2+}$ (A) and vascular (J) response traces evoked by whisker stimulation in PA, PSs, first- and second-order capillaries at D0 and at different post-stroke time points. Each curve represents the mean response overlaid from eight mice, with post-stroke days indicated by color.

(B−I) Amplitude of negative (B−E) and positive (F−I) $Ca^{2+}$ peaks in PAs, PSs, first- and second-order capillaries at D1, D4, D7, D14, and ≥ D21 post-stroke, normalized to baseline (D0). Data represent n = 8–16 ROIs from 8 mice in PAs, n = 6–10 ROIs from 8 mice in PSs, n = 8–15 ROIs from 8 mice in first-order capillaries, and n = 7–18 ROIs from 8 mice in second-order capillaries.

(K−N) Amplitude of peak vascular responses in PAs (K), PSs (L), first- (M) and second-order (N) capillaries at D1, D4, D7, D14, and ≥ D21 post-stroke, normalized to baseline (D0). Data represent n = 8–16 ROIs from 8 mice in PAs, n = 6–10 ROIs from 8 mice in PSs, n = 8–16 ROIs from 8 mice in first-order capillaries, and n = 15–26 ROIs from 8 mice in second-order capillaries.

Data are presented as mean ± SEM. Bayesian state-space analysis was applied for panels B−I and K−N. * indicates a statistically supported conclusion (posterior probability > 0.95) for comparisons between baseline and post-stroke time points.

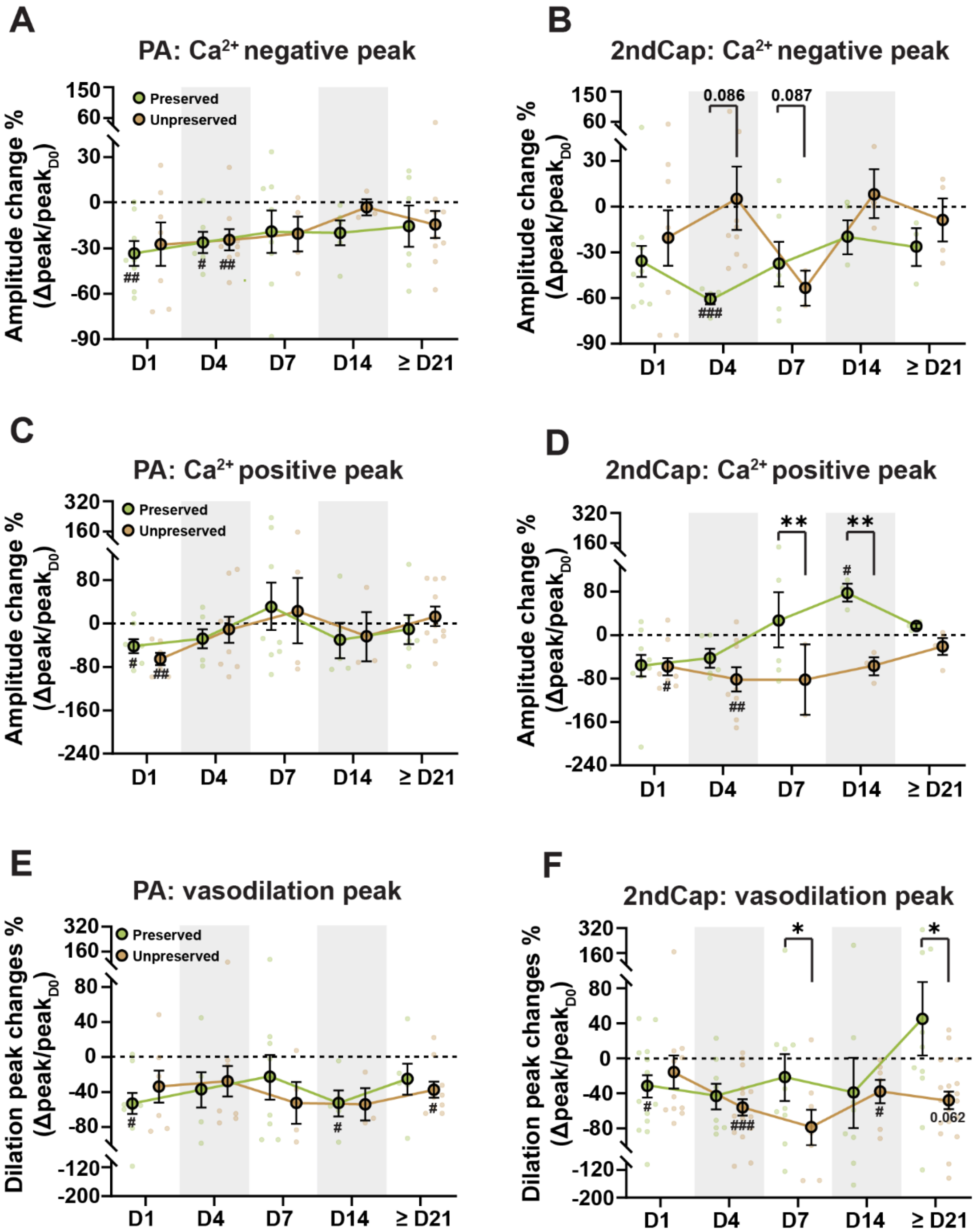


**Supplemental Figure 8. Ischemia induces differential reductions in mural cell calcium and vascular responses to whisker stimulation.**

(A−D) Amplitude of negative (A, B) and positive (C, D) $Ca^{2+}$ peaks in PAs, and second-order capillaries at D1, D4, D7, D14, and ≥ D21 post-stroke, normalized to baseline (D0). Green denotes the preserved group, and brown denotes the unpreserved group. Data represent n = 3–10 ROIs from 8 mice in PAs and n = 2–10 ROIs from 8 mice in second-order capillaries.

(E, F) Amplitude of peak vascular responses to whisker stimulation in PAs (E) and second-order capillaries (F) at D1, D4, D7, D14, and ≥ D21 post-stroke, normalized to baseline (D0). Green denotes the preserved group, and brown denotes the unpreserved group. Data represent n = 3–10 ROIs from 8 mice in PAs and n = 7–19 ROIs from 8 mice in second-order capillaries.

Data are presented as mean ± SEM. A linear mixed-effects model was applied for A−F. Comparisons between baseline and post-stroke time points are indicated by # $P < 0.05$, ## $P < 0.01$, and ### $P < 0.001$. Comparisons between preserved and unpreserved groups are indicated by * $P < 0.05$, and ** $P < 0.01$.

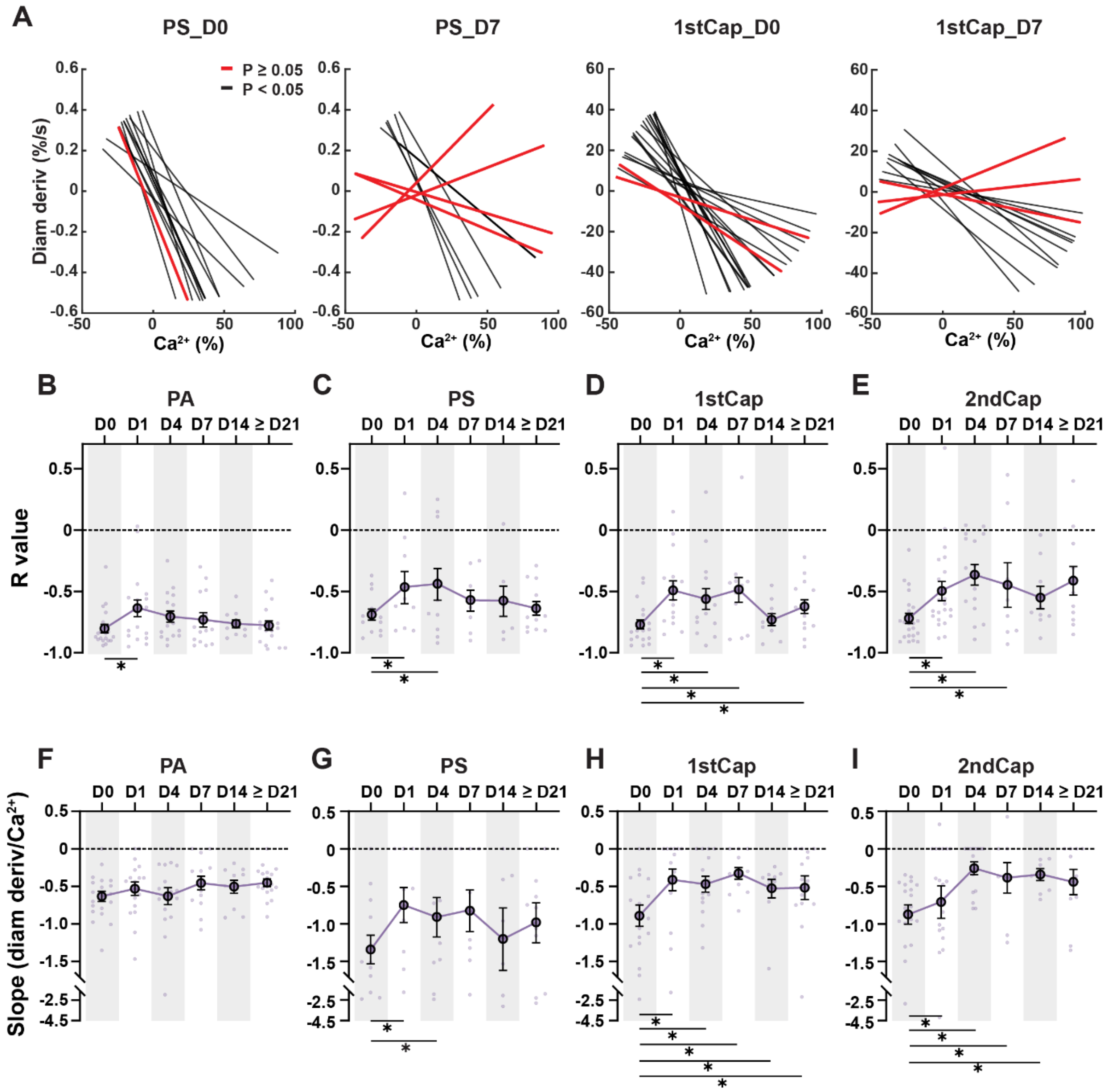


**Supplemental Figure 9. Ischemia induces uncoupling between calcium dynamics and vascular responses.** (A) Regression lines illustrating correlations between mural cell $Ca^{2+}$ signals and vascular responses in PSs and first-order capillaries at D0 and D7 across eight mice. Black lines indicate statistically significant correlations ($P < 0.05$), whereas red lines indicate non-significant correlations ($P \geq 0.05$).

(B−I) Summary of Pearson correlation coefficients (R; B−E) and slopes of regression lines (F−I) describing the relationship between $Ca^{2+}$ signals and the derivative of vascular diameter in PAs, PSs, first- and second-order capillaries at D0 and various days post stroke. Data represent n = 10–20 ROIs from 8 mice in PAs, n = 7–13 ROIs from 8 mice in PSs, n = 10–19 ROIs from 8 mice in first-order capillaries, and n = 8–22 ROIs from 8 mice in second-order capillaries.

Data are presented as mean ± SEM. Bayesian state-space analysis was applied for panel B−I. * denotes a statistically supported conclusion (posterior probability >0.95) for comparisons between baseline and post-stroke time points.

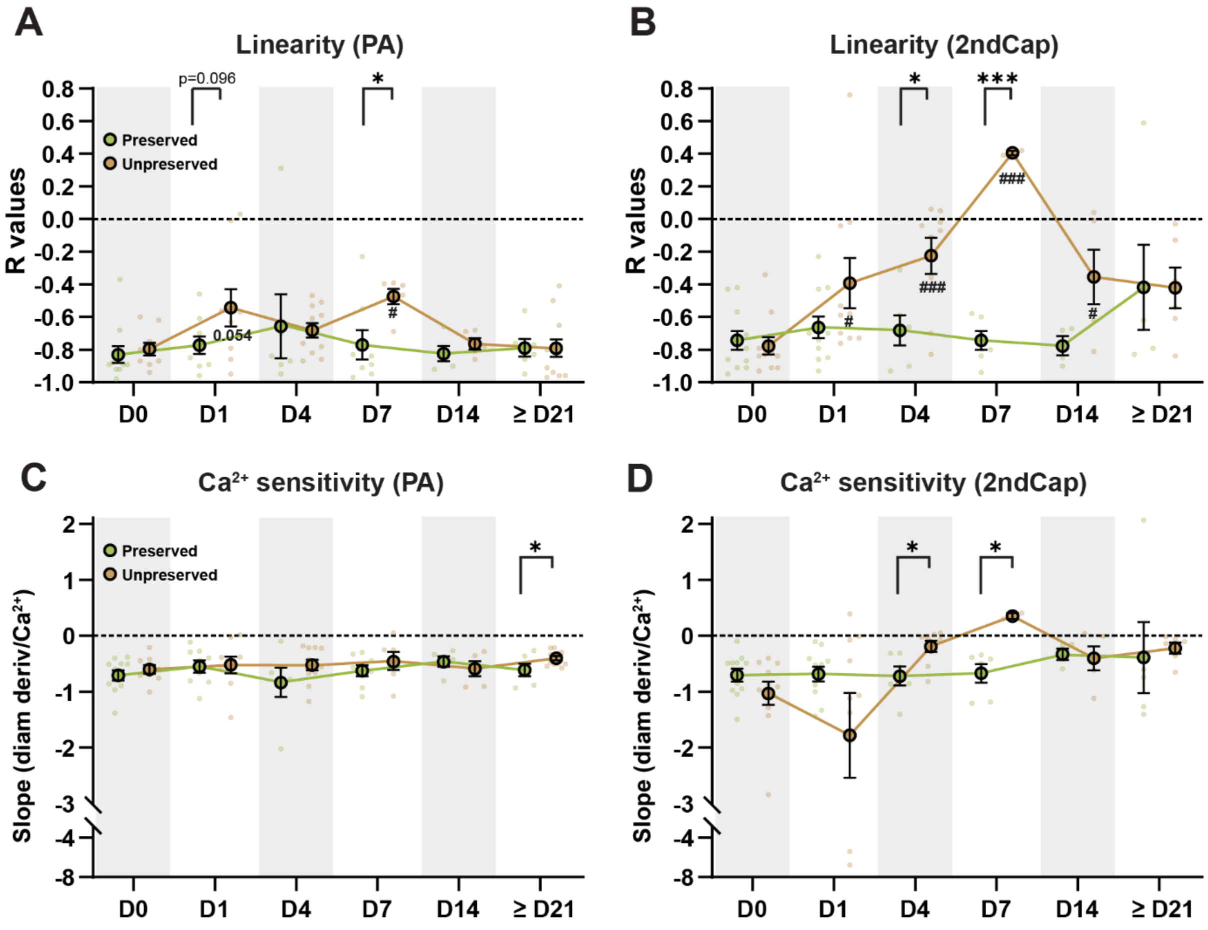


**Supplemental Figure 10 Ischemia induces uncoupling between calcium dynamics and vascular responses, particularly in the unpreserved group.**

(A, B) Summary of Pearson correlation coefficients (*R*) for correlations between $Ca^{2+}$ signals and the derivative of vascular diameter in PAs (A) and second-order capillaries (B) at D0 and various days post stroke. Data represent n = 5–11 ROIs from 8 mice in PAs and n = 2–11 ROIs from 8 mice in second-order capillaries. Green denotes the preserved group, and brown denotes the unpreserved group.

(C, D) Slopes of the regression lines describing the relationship between $Ca^{2+}$ signals and the derivative of vascular diameter in PAs (C) and second-order capillaries (D) at D0 and various days post stroke (n = 5−11 ROIs from 8 mice in PAs; n = 2−11 ROIs from 8 mice in second-order capillaries). Green denotes the preserved group, and brown denotes the unpreserved group.

Data are presented as mean ± SEM. A linear mixed-effects model was applied for A−D. Comparisons between baseline (D0) and post-stroke time points are indicated by # $P < 0.05$, ## $P < 0.01$, and ### $P < 0.001$. Comparisons between preserved and unpreserved groups are indicated by * $P < 0.05$, ** $P < 0.01$, and *** $P < 0.001$.
